# Lipin-1 restrains macrophage lipid synthesis to promote inflammation resolution

**DOI:** 10.1101/2023.10.23.563587

**Authors:** Temitayo T. Bamgbose, Robert M. Schilke, Oluwakemi O. Igiehon, Ebubechukwu H. Nkadi, David Custis, Sushma Bharrhan, Benjamin Schwarz, Eric Bohrnsen, Catharine M. Bosio, Rona S. Scott, Arif Yurdagul, Brian N. Finck, Matthew D. Woolard

## Abstract

Macrophages are critical to maintaining and restoring tissue homeostasis during inflammation. The lipid metabolic state of macrophages influences their function, but a deeper understanding of how lipid metabolism is regulated in pro-resolving macrophage responses is needed. Lipin-1 is a phosphatidic acid phosphatase with a transcriptional coregulatory activity (TC) that regulates lipid metabolism. We previously demonstrated that lipin-1 supports pro-resolving macrophage responses, and here, myeloid-associated lipin-1 is required for inflammation resolution, yet how lipin-1-regulated cellular mechanisms promote macrophage pro-resolution responses is unknown. We demonstrated that the loss of lipin-1 in macrophages led to increased free fatty acid, neutral lipid, and ceramide content and increased phosphorylation of acetyl-CoA carboxylase. The inhibition of the first step of lipid synthesis and transport of citrate from the mitochondria in macrophages reduced lipid content and restored efferocytosis and inflammation resolution in lipin-1^m^KO macrophages and mice. Our findings suggest macrophage-associated lipin-1 restrains lipid synthesis, promoting pro-resolving macrophage function in response to pro-resolving stimuli.

**Teaser:** Lipin 1 blockade of lipid biosynthesis inducing mitochondrial citrate export promotes efferocytosis and inflammation resolution.

## Introduction

Acute inflammation in response to noxious stimuli, often a beneficial response, must be resolved (*1*). Resolution of inflammation is an active process that facilitates disease resolution and prevents the harmful effect of unresolved inflammation (*2*). An impairment in the resolution of inflammation is implicated in the pathophysiology of cardiometabolic and chronic inflammation-driven diseases (*3*). Macrophages are critical to initiating and resolving inflammatory responses (*4*, *5*). To effectively carry out both inflammation and inflammation resolution, macrophages must polarize toward distinct phenotypes characterized by drastic transcriptional, translational, and metabolic changes (*6*, *7*). Dysregulation in one or more of these pathways can significantly impact the macrophages’ ability to function appropriately.

Pro-inflammatory macrophages quickly upregulate glycolysis to meet energy demands and carry out inflammatory or antimicrobial responses such as cytokine production and phagocytosis (*8–10*). Though glycolysis-derived pyruvate feeds into the tricarboxylic acid cycle (TCA) cycle in pro-inflammatory macrophages, the deactivation of succinate dehydrogenase and isocitrate dehydrogenase disrupts oxidative metabolism, leading to the accumulation of citrate (*10*, *11*). Citrate can be exported from the mitochondria by the mitochondrial citrate carrier (CIC)/citrate transport protein (CTP) to generate oxaloacetate-derived NAPDH for ROS production and acetyl-CoA for de novo fatty acid biosynthesis (*10–12*). Inhibition of the succinate dehydrogenase (SDH) complex leads to a buildup of succinate, promoting inflammatory cytokine production (*11*, *13*, *14*). Since the TCA cycle generates electron carriers and the SDH is part of complex II of the electron transport chain, a dysfunctional TCA cycle impairs oxidative phosphorylation. Pro-resolving macrophages initially utilize glycolysis before switching to fatty acid oxidation for metabolic needs. Though glycolysis is dispensable for pro-resolving polarization (*15*), it is suggested that glycolysis can fuel fatty acid biosynthesis to provide substrates for fatty acid oxidation in pro-resolving macrophages (*11*). Unlike pro-inflammatory macrophages, pro-resolving macrophages have a functional TCA cycle that supports oxidative phosphorylation (*10*, *11*)

Free fatty acid metabolism can regulate macrophage polarization and effector functions (*16*). Accumulation of fatty acid-derived phospholipids and triglycerides is a feature of LPS-activated macrophages as de novo fatty acid synthesis supports pro-inflammatory macrophage activation, recruitment, phagocytosis, and production of inflammatory cytokines (*16*, *17*). The contribution of lipid metabolism to pro-resolution responses is less well-defined and more controversial. IL-4 stimulation initially leads to increased beta-oxidation but eventually requires SREBP-mediated de novo lipid synthesis to restore lipid pools (*18*, *19*). Even the contribution of beta-oxidation is controversial. While beta-oxidation does not appear to be required for canonical M2 gene expression (*20*), beta-oxidation may promote pro-resolving macrophage effector functions and disease resolution (*21–23*). Specifically, the enhancement of fatty acid oxidation and oxidative phosphorylation enhanced the uptake of necroptotic cells (*22*). Also, the catabolism of Apoptotic cell (AC)-derived lipids orchestrates the production of IL-10, which promotes tissue repair (*21*). Altogether, one must wonder about the upstream molecular events that decide the fate of fatty acids and how their utilization aligns with pro-resolving macrophage responses.

Lipin-1 is a phosphatidic acid phosphatase that converts phosphatidic acid into diacylglycerol (*24*). Mutations in the gene encoding lipin-1 (LPIN1) lead to several metabolic syndromes in mice (*25*) and severe, sometimes fatal, episodic rhabdomyolysis in people (*26–28*). Loss of lipin-1 also results in cardiac lipotoxicity (*29*, *30*), mitochondria dysfunction (*29*), insulin resistance (*31*), and atherosclerosis (*32*). In macrophages, the enzymatic activity of lipin-1 supports a pro-inflammatory response (*33*, *34*). Lipin-1 also has an enzymatically independent transcriptional coregulator activity in which lipin-1 binds to transcription factors, such as Peroxisome proliferator-activated receptors (PPARs) and Sterol regulatory element binding proteins (SREBPs), to regulate their activity and subsequent gene expression (*35*). In non-myeloid cells, lipin-1 augmentation of PPAR activity upregulates FA utilization while suppressing lipid synthesis by inhibiting SREBP (*35*, *36*). Altogether, during processes that require lipid catabolism, lipin-1 has been shown to suppress FA synthesis and upregulate lipid catabolism preferentially (*23*, *35*, *37*). Previously, we demonstrated that lipin-1 non-enzymatic activity promotes pro-resolving macrophage polarization, β-oxidation, and efferocytosis (*23*, *38*). We have also shown that in response to pro-resolving stimuli, loss of lipin-1 leads to a build-up of metabolites that may contribute to lipid synthesis (*23*). This data suggests that lipin-1 potentially regulates lipid metabolism to align with macrophage function. However, the aspect of lipin-1-regulated lipid metabolism that supports pro-resolving macrophage function is unknown.

In this study, we have used transgenic mouse models lacking the enzymatic activity of lipin-1 or both activities of lipin-1 in myeloid cells to report that lipin-1 non-enzymatic activity promotes disease resolution and pro-resolving macrophage responses by restraining fatty acid biosynthesis.

## Results

### Myeloid-associated lipin-1 promotes inflammation resolution

We have previously demonstrated that lipin-1 contributes to macrophage efferocytosis (*23*), and deletion of myeloid-associated lipin-1 delayed full skin wound closure (*38*) and increased atherosclerosis progression (*32*). Collectively, our previous studies suggest that lipin-1 is involved in inflammation resolution. We used a zymosan-induced peritonitis inflammation model (Fig 1A) to investigate the contribution of myeloid-associated lipin-1 to the resolution of inflammation. Mice lacking lipin-1 in myeloid cells (lipin-1^m^KO) and their littermate controls were intraperitoneally (IP) injected with 0.1mg zymosan to induce peritonitis. Clearance of neutrophils after the onset of zymosan-induced inflammation is a marker for inflammation resolution (*5*). We quantified neutrophils (CD45+ CD11b+ Ly6g+) and monocytes/macrophages (CD45+ CD11b+ Ly6g- F4/80+) numbers by collecting peritoneal cells by lavage, staining them, and performing flow cytometry (Fig S1A). Consistent with previous reports using this model (*39*, *40*), we observed a rapid influx of leukocytes after 4 hours of injection (Fig 1B-D), with the number of neutrophils peaking at 12 hours after injection (Fig 1C). There was a delay in the clearance of neutrophils within the lipin-1^m^KO mice compared to wild-type mice, suggesting a defect in inflammation resolution with 10 hours increase in resolution index compared to WT mice (Fig. 1D). We observed no significant difference in the number of recruited monocytes/macrophages, meaning the difference in PMNs is due to clearance and not the presence of higher numbers of macrophages (Fig 1E). We further examined if a difference in the number of polymorphonuclear leukocytes (PMNs) was observed in lipin-1 enzymatic KO mice. In contrast, lipin-1 enzymatic KO mice are not impaired in their ability to resolve inflammation (Fig. S1B-D), suggesting that the observed phenotype was independent of the enzymatic activity.

**Fig. 1.**
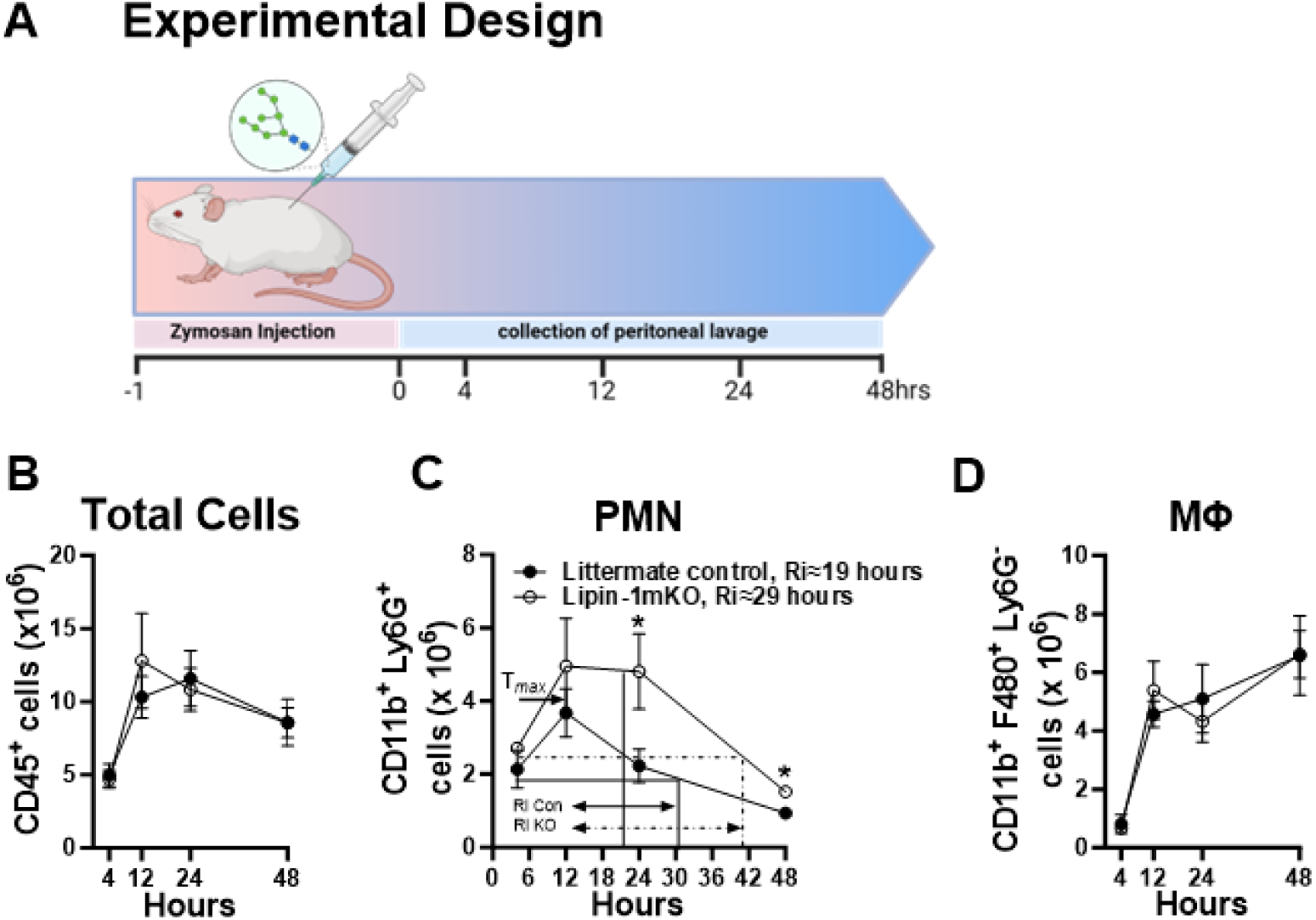
Loss of myeloid lipin-1 delays inflammation resolution. (**A**) Mice were subjected to a zymosan challenge (0.1mg/mouse). PMNs and macrophages were quantified from the peritoneal cavity by flow cytometry. (**B**) Total number of cells isolated from the peritoneal cavity. (**C**) Total number of PMNs isolated from the peritoneal cavity. Resolution interval (Ri) was determined. (**D**) Total number of Macrophages isolated from the peritoneal cavity. Illustration in (**A**) was created using BioRender.com. A minimum of three mice per group per time point were analyzed. N ≥3 mice per group per time point. Values are means ± SEM. Unpaired two-tailed T-tests were performed between groups at each time point. *= p≤0.05.

### Lipin-1 promotes a pro-resolution metabolic phenotype and attenuates inflammation in vivo

The metabolic status of macrophages is critical to their ability to perform proper efferocytosis, which is crucial for inflammation resolution (*21*, *23*, *40*). We have previously demonstrated *in vitro* that the loss of lipin-1 increased glycolysis, broke the TCA cycle with increased citrate and isocitrate, and reduced oxidative phosphorylation (*23*). We sought to determine if similar metabolic alterations were seen *in vivo* during inflammation resolution. We performed a high-dimensional, flow cytometric analysis of immune cell populations and metabolic targets that correlated with the metabolic state of the cells isolated from the peritoneal cavity during Zymosan-induced peritonitis (*41*). We examined macrophage phenotypes six days after injection of 0.1mg zymosan, which correlates with peak pro-resolution macrophage response (*42*). We isolated cells from the peritoneal cavity by lavage (*41*). See Table S1 for antibody targets. No significant change in macrophage (Lin^-^ CD11b^+^ F4/80^+^ CD45^+^) percentage or number was observed (Fig 2A). In lipin-1^m^KO peritoneal macrophages (pMACs), there was a significant increase in the Median Fluorescent Intensity (MFI) of Glut1 (Fig 2B), suggesting a potential glycolysis increase supported by our previous study (*23*). We also report a significant decrease in MFI of Cytochrome C (CytC) and succinate dehydrogenase (SDHA), components of the electron transport chain and Krebs cycle (Figure 2B) in lipin-1 deficient pMACs (Fig 2B). Low levels of CytC and SDHA suggest reduced mitochondria content, impaired mitochondria function, and a defective Krebs cycle. These data are suggestive of impaired oxidative metabolic capacity, a phenotype we previously observed of lipin-1 KO macrophages *in vitro* (*23*). We observe a significant decrease in Siglec F and an increase in Ly6C and MHCII staining, suggesting a more pro-inflammatory state of macrophages from lipin-1^m^KO mice (Fig 2C).

**Fig. 2.**
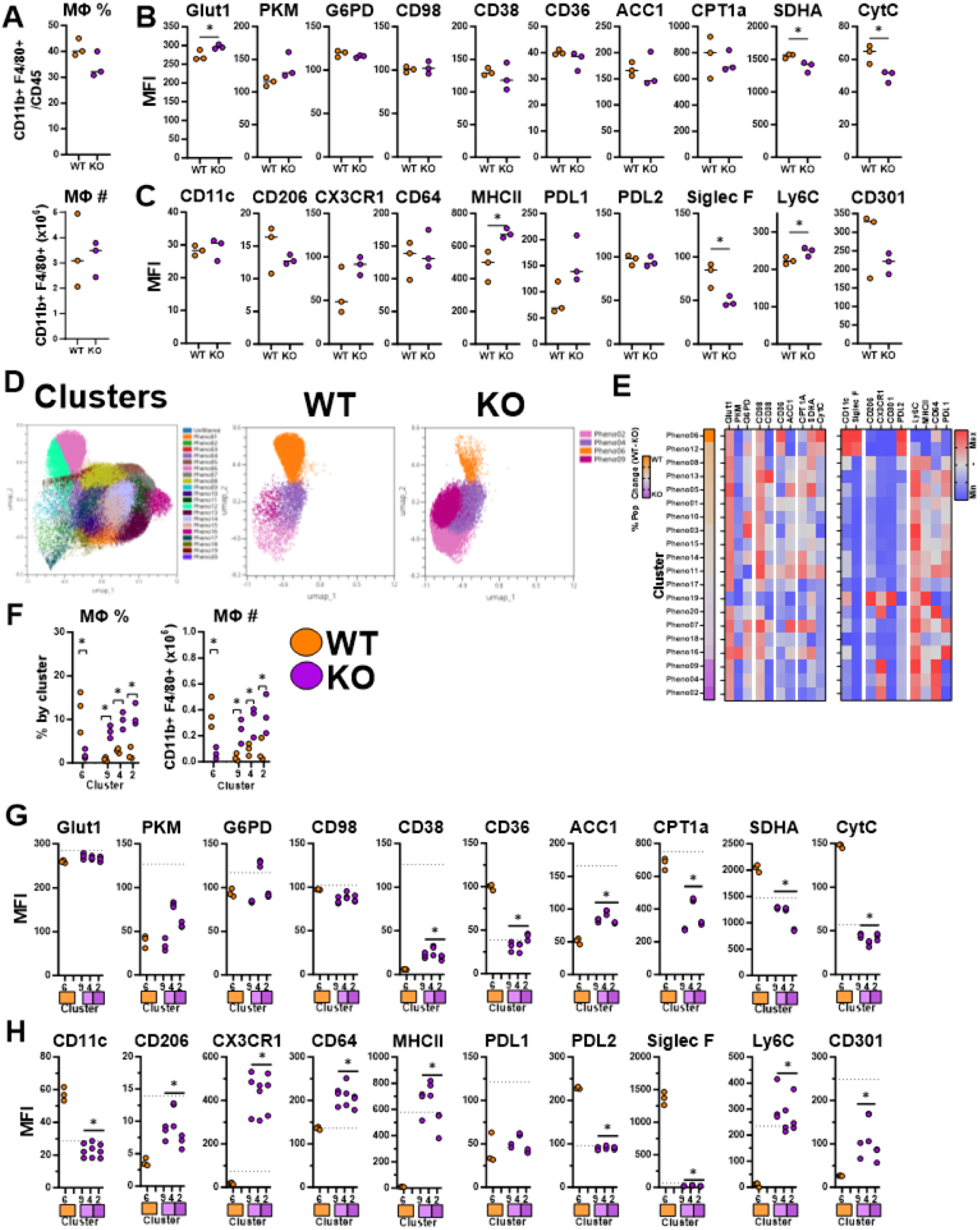
Lipin-1 promotes pro-resolution phenotype in vivo. Mice were challenged with 0.1mg zymosan, and six days later, peritoneal cells were isolated by lavage. Isolated cells were stained for flow cytometric analysis. (**A**) The number and ratio of CD11b+ F4/80+ lin- cells in the peritoneal cavity. (**B-C**) Median fluorescent Intensity of metabolic and inflammatory markers of CD11b+ F4/80+ lin- cells. (**D**) UMAP and phenograph clustering of macrophages and those enriched in wild-type (orange) and lipin-1mKO (purple) mice using metabolic markers. (**E**) heat map of Percent population changes of phenoclusters between wild-type and lipin-1mKO mice and Min-Max heat map of metabolic and inflammatory markers from identified phenoclusters. (**F**) The number and ratio of CD11b+ F4/80+ lin- cells by phenocluster type. (**G-H**) Median fluorescent Intensity of metabolic and inflammatory markers of CD11b+ F4/80+ lin- cells from phenocluster 6 in wild-type mice (orange) and phenoclusters 9, 4, & 2 from lipin-1^m^KO (purple) mice. N=3 mice per group. Dots represent individual mice; lines are means. Unpaired Two-tailed T-test was performed between groups to define significance *= p≤0.05 Abbreviations: **Glut 1**: glucose transporter, **PKM**: Pyruvate Kinase M2, **G6PD**: Glucose-6- phosphate dehydrogenase, **ACC1**: Acetyl CoA carboxylase 1, **CPT1A**: mitochondria fatty acid importer, **SDHA**: Succinate dehydrogenase, **CytC**: Cytochrome C, **PDL**: Programmed Death-Ligand

*In vivo*, macrophages are heterogeneous in metabolic and inflammatory phenotypes (*41*). As such, we examined this heterogeneity by dimensional reduction and clustering of concatenated samples of Lin^-^ CD11b+ F4/80^+^ CD45 cells, concentrating on metabolic markers identified in Table S1. A total of 20 clusters were defined, and clusters that predominantly contained WT or KO were identified (Fig 2D-F) and used for further analysis. Cluster 6 was primarily comprised of WT macrophages, while clusters 9, 4, and 2 were predominantly comprised of lipin-1 KO macrophages (Fig 2D-F). Analysis of these clusters for their expression of metabolic markers showed that loss of lipin-1 leads to a decrease in CPT1A protein, a reduction in surface expression of CD36, a scavenger receptor known to mediate apoptotic cell clearance, and a significant decrease in Cytochrome C, and SDHA, which are consistent with the global pMACs staining (Fig 2G). CPT1a controls the first committed step of beta-oxidation, suggesting these macrophages may have reduced beta-oxidation capacity, a phenotype we observe *in vitro* (*23*). Additionally, the decreased Cytochrome C and SDHA would indicate defective oxygen consumption and could lead to increased citrate and isocitrate, observations we have seen of *in vitro* lipin-1^m^KO macrophages stimulated with IL-4 (*23*). We also observe an increase in ACC in lipin-1 KO pMACs, though below the total cluster average, suggesting that loss of lipin-1 leads to an increase in fatty acid (FA) biosynthesis in a specific population of macrophages. In sum, these metabolic phenotypes of macrophages *in vivo* align well with our previous *in vitro* data.

We also looked at the inflammatory profile of the pMACs within these clusters. Our data suggests that there were more myeloid-derived inflammatory pMACs with an observed increase in MHCII, PDL2, Ly6C, CX3CR1, and CD64, the high-affinity IgG FC receptor (Fig 2H). An increase in CX3CR1 and Ly6C, known to be high in bone marrow monocytes (43, 44), suggests ongoing inflammation and a high number of inflammatory macrophages recently generated from recruited inflammatory monocytes (Fig 2H). Lipin-1-dependent increase in PDL2, which inhibits T cell proliferation and cytokine production (*45*, *46*), suggests that lipin-1 promotes inflammation resolution and further supports the notion of ongoing inflammation in the lipin-1^m^KO mice (Fig 2H). In addition, the increase in surface MHCII expression suggests not only unresolved inflammation but also defective efferocytosis since efficient efferocytosis prevents the antigen presentation of efferocytotic cargo peptides, and one such mechanism is via the degradation of efferocytotic cargo peptides in such a manner that they become unsuitable for antigen presentation by MHC class II (*3*, *47*).

Interestingly, though below the total cluster average, we observe a significant increase in CD206 and CD301, which are generally classified as M2 Macrophage markers in lipin-1^m^KO pMAC clusters. We hypothesize that at the 6-day time point, where WT pMACs might have returned to a ground state, lipin-1 KO pMACs might still be expressing M2 markers due to a delay in polarization to the pro-resolving phenotype. The result from the lipin-1^m^EnzyKO suggests that these phenotypic changes in metabolism and inflammatory profile are independent of the enzymatic activity of lipin-1 as we observe no differences in their metabolic and inflammatory markers staining compared to WT (Fig. S2). Collectively, our data align with what we have previously demonstrated *in vitro*, suggesting that the metabolic defects observed in lipin-1 deficient macrophages are likely responsible for the defects in inflammation resolution.

### Lipin-1 regulates the production and channeling of lipids

Lipin-1 regulates lipid synthesis in several other cell types (*35*, *36*), and both our previous (*23*) and current data (Fig. 2G) suggest that lipin-1 restrains lipid synthesis in macrophages in response to pro-resolving stimuli. We had previously demonstrated elevated citrate (*23*), a precursor for fatty acid synthesis, in lipin-1 deficient macrophages. Flow cytometric analysis of *in vivo* macrophages demonstrated increased ACC1 and decreased CPT1a protein levels (Fig. 2G), suggesting that loss of lipin-1 may promote lipid accumulation in macrophages during pro-resolving responses. We investigated the distribution and composition of lipids in IL-4 stimulated Bone marrow marrow-derived macrophages (BMDMs) from lipin-1^m^KO mice and littermate controls by liquid chromatography-tandem spectrometry (LC–MS/MS). Principal component analysis (PCA) from our lipidomics assay showed a distinct difference in the lipid profile of WT and Lipin-1^m^KO BMDMs under unstimulated conditions that were largely maintained or exacerbated when stimulated with IL-4 (Fig. 3A). When lipid species are grouped by family, we observe alterations in the lipid composition of lipin-1 KO macrophages (Fig. 3B-E). There was an increase in fatty acid levels at both baseline and in response to IL-4 in the lipin-1 KO macrophages (Fig. 3B-C, 3E). Though our data does not demonstrate an IL-4-dependent increase in fatty acid levels in macrophages deficient in lipin-1, the observed IL-4-dependent increase in the levels of triglycerides (TAG), diacylglycerol (DAG), cholesterol esters (CE), and several phospholipids (Fig 3B-C, 3E) supports that excess fatty acids accumulating in response to IL-4 in lipin-1 KO BMDMs are being channeled for storage. We also confirmed this buildup of lipid storage species by Nile red staining (Fig 3F). In addition to lipid storage in triglycerides, sphingolipid biosynthesis responds to excess free fatty acids. The buildup of bioactive sphingolipids such as ceramides induces cellular dysfunction and impairs pro-resolving macrophage function (*48–51*). Our lipidomic analysis of macrophages deficient in lipin-1 shows a trending rise in total ceramides and an IL-4-dependent increase in dihydroceramide, suggesting an increase in de novo sphingolipid biosynthesis (Fig 3B-E).

**Fig. 3.**
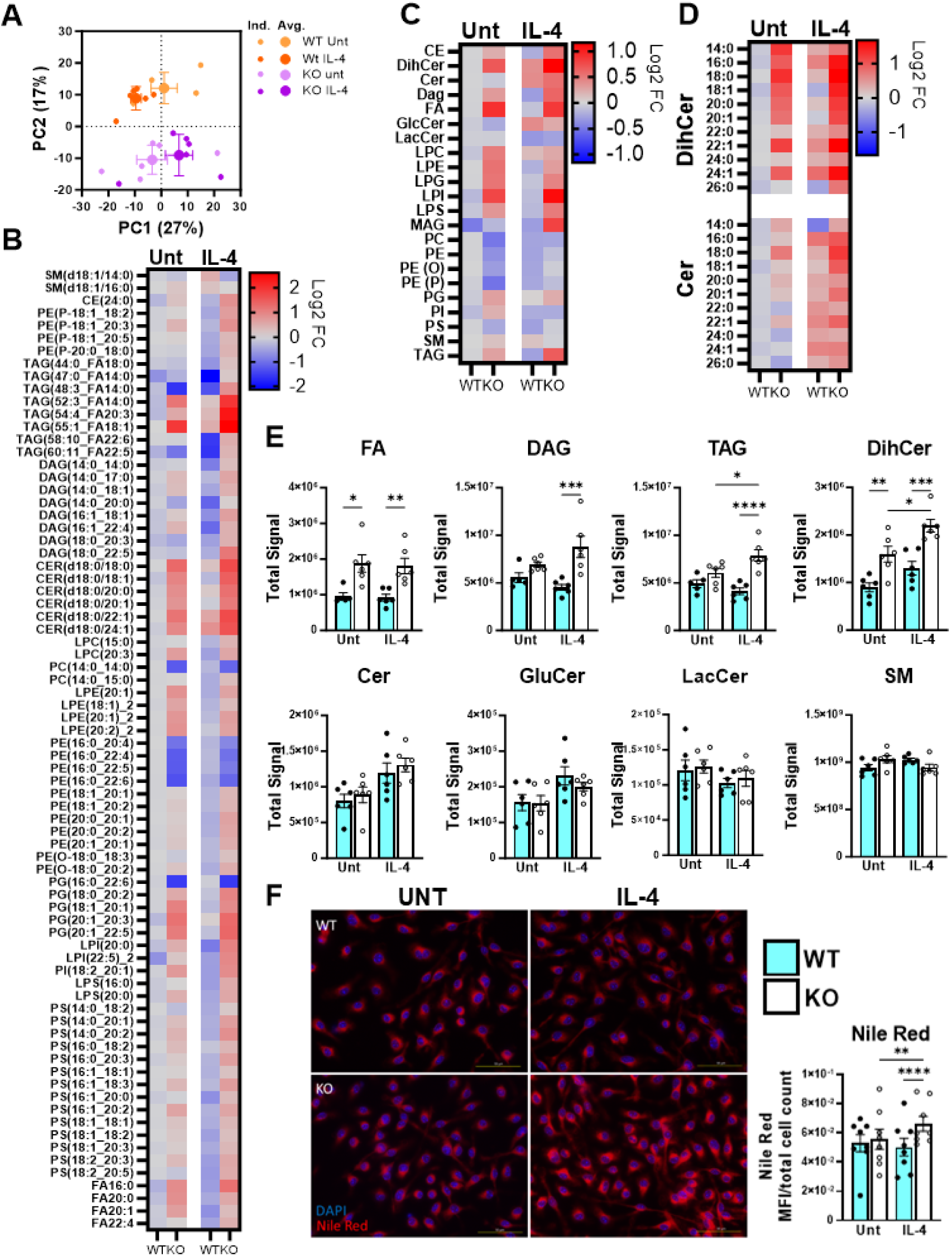
Loss of lipin-1 results in an increase in fatty acids, neutral lipids, and ceramides. Lipids harvested from IL-4-stimulated (40ng/ml, 4 hours) BMDMs were processed via LC–MS for lipidomics analysis. (**A**) PCA analysis of z-score values shows the spatial distribution of lipids within each condition. (**B**) Heat map of most-changed lipid species relative to untreated cells (n=6). A 3% FDR correction of student’s t-test analysis between fold change adjusted IL-4 stimulated WT and KO values was used to populate lipids in panel B (**C**) Heat map of lipid species as a family (n=6). Lipids that are represented by family in (C) include all respective species that passed the 3000 units cut-off, regardless of statistical significance. (**D**) Quantified signals of lipid species from the ceramide and dihydroceramide family (n=6). (**E**) Quantification of lipid families in C.(**F**) Nile red staining images of IL-4 stimulated (40ng/ml, 4 hours) BMDMs. Heatmaps were made of Log2 Fold Change Analysis from WT untreated. Significance was determined by one-way ANOVA *= p≤0.05, **= p≤0.01, ***= p≤0.001, and ****= p≤0.001. Lipid abbreviations: **FA**: Fatty acids, **DAG**: Diglycerides, **TAG**: Triglycerides, **DihCer**: Dihydroceramide, **Cer**: Ceramide, **GlcCer** Glucosylceramide**, SM:** Sphingomyelin, **MAG**: Monoacylglycerol, **LPC:** Lysophosphatidylcholine, **LPE:** Lysophosphatidylethanolamine, **LPI**: Lysophosphatidylinositol, **LPS**: Lysophosphatidylserine, **LPG**: Phosphatidylglycerol, **PE**: Phosphatidylethanolamine, **PC**: Phosphatidylcholines, **PG**: Phosphatidylglycerol, **PS**: Phosphatidylserine, **PI**: Phosphatidylinositol, **PE O**: Ether-linked Phosphatidylethanolamine, **PE P**: Phosphatidylethanolamine plasmalogen.

### LIPIN-1 RESTRAINS FA SYNTHESIS IN RESPONSE TO PRO-RESOLVING STIMULI

The build-up of fatty acids and neutral lipids observed in IL-4-stimulated lipin-1 KO macrophages might result from increased lipid uptake, reduced lipid catabolism, or de novo fatty acid synthesis (FIG 4A). Though we have previously shown that macrophage lipid uptake is independent of lipin-1 (*23*), the loss of lipin-1 could be causing an increase in the expression of enzymes involved in fatty acid biosynthesis or activation of fatty acids. We quantified the amount of these enzymes by western blot. We observe no difference in Cytoplasmic acetyl-CoA synthetase (AceCS1) protein levels at baseline and in response to IL-4 (FIG 4B). Citrate is the first metabolite in free fatty acid biosynthesis, where it is transported from the mitochondria by the citrate transport protein (CTP) or citrate carrier (CIC) (FIG 4A). We observe no difference in the protein levels of CTP, and we also do not see differences in the protein levels of fatty acid synthase (FAS), which catalyzes palmitate biosynthesis from malonyl-CoA (Fig 4B). These observations suggest that lipid biosynthesis may be upregulated without significantly increasing enzyme abundance, potentially by post-translational regulatory mechanism. Alternatively, basal abundance of these enzymes might be sufficient to handle increasing inputs of fatty acid biosynthesis metabolites.

**Fig. 4.**
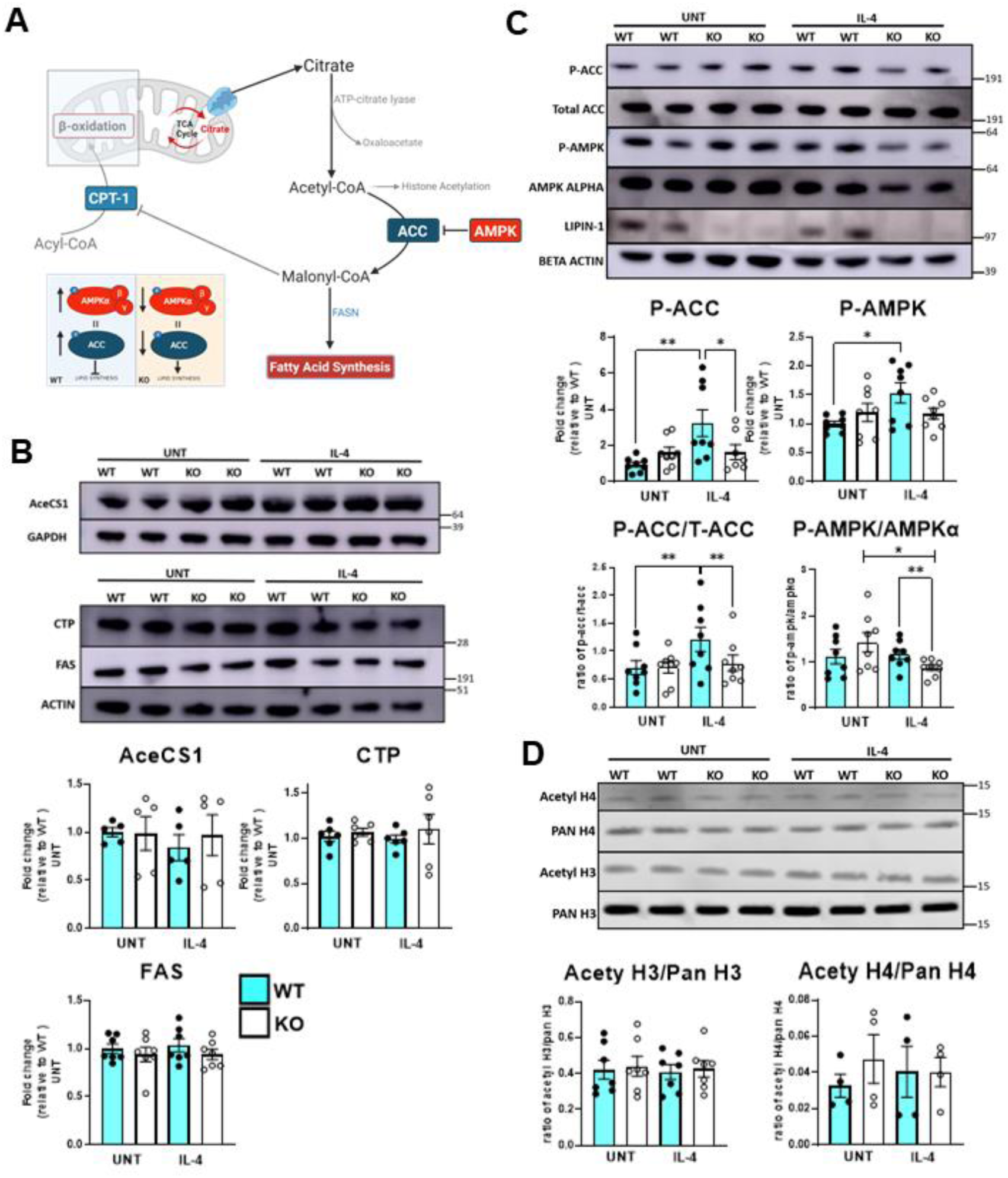
Loss of lipin-1 during IL-4 stimulation results in a failure to restrain fatty acid biosynthesis. (**A**) Illustration of the de novo fatty acid biosynthesis pathway (**B-C**) Protein was isolated from IL-4-stimulated (40ng/ml, 4 hours) whole cells, and protein abundance of fatty acid biosynthesis enzymes and proteins was quantified by Western blot analysis (n>3). (**D**) Representative western blot images and quantified protein abundance of histone proteins and post-translational modifications (n=X). Illustration in (**A**) was created using BioRender.com. Bars represent the standard error of the mean (± SEM). Significance was determined by one-way ANOVA (fold change) and paired student’s t-test (ratio). *= p≤0.05, **= p≤0.01.

Acetyl-CoA carboxylase (ACC) conversion of acetyl-CoA to malonyl-CoA is a rate-limiting step of fatty acid synthesis. ACC is negatively regulated by AMP-activated protein kinase (AMPK) phosphorylation at serine 79 (*52*). In response to IL-4, phosphorylation of AMPK and ACC is increased in wild-type BMDMs but not in lipin-1 KO BMDMs (Fig 4C). We also observed similar ACC phosphorylation results when using apoptotic cells as an alternative pro-resolving stimulus (Fig S3A). Lipin-1 KO BMDMs are missing both lipin-1 activities; we performed the same experiment, using BMDMs from KO mice lacking enzymatic activity but with preserved transcriptional regulatory function to determine which lipin-1 activity was responsible for the observed phenotype. We observed an increase in ACC and AMPK phosphorylation in both WT and Lipin-1 EKO macrophages, suggesting that lipin-1 non-enzymatic activity is responsible for the decreased phosphorylation of ACC and AMPK in lipin-1^m^KO macrophages (Fig S3B).

Previously, we reported an IL-4-dependent decrease in Acetyl CoA in lipin-1 KO BMDMs (*23*). Acetyl CoA can be used in numerous processes, including de novo FA biosynthesis or epigenetic modifications, which contribute significantly to macrophage biology and function (*16*). We propose that the previously reported elevated citrate leads to increased production of acetyl-CoA, which is utilized for lipid synthesis. However, it is possible that acetyl-CoA is being used up for epigenetic modifications. We report no differences in histone acetylation between WT and lipin-1 deficient macrophages (Fig 4D). This further supports a more causative effect of increased acetyl CoA utilization for FA synthesis. Collectively, loss of lipin-1 does not necessarily lead to an upregulation of the FA biosynthesis pathway but a failure to restrain FA synthesis at the citrate carrier and ACC checkpoints.

### Inhibition of the citrate carrier restores efferocytic function

We previously demonstrated that lipin-1 was required for efficient efferocytosis (*23*), but only after macrophages encountered the first apoptotic cell. IL-4 can enhance macrophage efferocytosis (*38*). We have shown that lipin-1 contributes to numerous IL-4-mediated responses in macrophages, including increased beta-oxidation, fatty acid metabolism, and pro-resolving gene expression (*23*, *38*). We aimed to investigate if IL-4 could produce a similar efferocytosis defect in lipin-1 KO BMDMs. After 6 hours of IL-4 treatment, we observed a defect in apoptotic cell uptake by lipin-1 KO BMDMs (Fig 5A), consistent with our previous data on phagocytosis of zymosan particles (*38*). These results suggest that lipin-1 is required for augmenting efferocytic capacity during pro-resolving elicited responses.

**Fig. 5.**
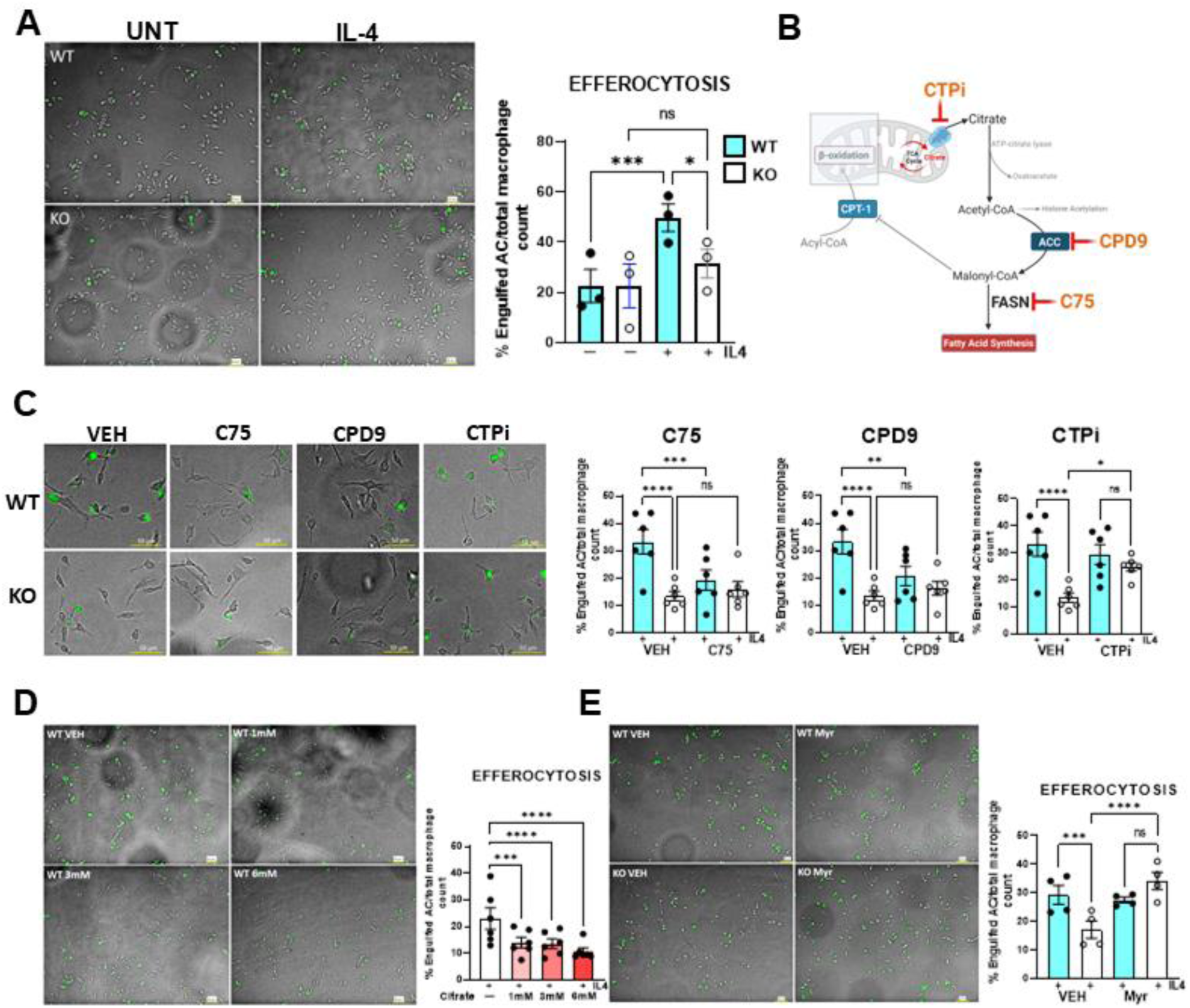
Inhibition of the citrate carrier and de novo ceramide biosynthesis restores efferocytic capacity in lipin-1 ko macrophages. (**A**) BMDMs from lipin-1^m^KO mice and littermate controls were stimulated with IL-4 for 6 hours and then challenged with CFSE-labeled apoptotic cells. (**B**) Illustration of the de novo fatty acid biosynthesis pathway and checkpoints of inhibition. (**C**) BMDMs were treated with 10µM inhibitors (FAS: *C75*, ACC: *Cpd9*, CTP: *CTPi*) and DMSO Vehicle (VEH) for 12 hours and subsequently cotreated with 40ng/ml IL4 and 10µM of respective inhibitors for 6 hours. Images were taken at 20x and zoomed at 1.5x to generate representative images. The experiment was done twice, each time with 3 unique pairs of individual WT and FKO (full KO) BMDMs (n= 6). At least 3 random images of each group were taken, quantified (**C**), and grouped to give individual dots. Individual experiments incorporated all inhibitors, so each inhibitor group had the same vehicle control. (**D)** BMDMs were treated with increasing concentrations of SCT (1mM, 3mM, 6mM) for 2 hours before co-treatment with IL-4 for 6 hours. In vitro efferocytosis was carried out (**E**). In vitro efferocytosis with 10µM Myriocin. BMDMs were treated with Myriocin for 2 hours and cotreated with Myriocin and IL-4 for 6 hours before the AC challenge. Illustration in (**B**) was created using BioRender.com. Bars represent the standard error of the mean (± SEM). Significance was determined by two-way ANOVA. *= p≤0.05, **= p≤0.01, ***= p≤0.001, and ****= p≤0.001.

We sought to determine whether alterations in free fatty acid metabolism in lipin-1 KO BMDMs were responsible for inhibiting efferocytosis capacity. Acetyl-CoA is the precursor for all free fatty acid synthesis (Fig 4A). Acetyl-CoA is converted into malonyl-CoA by ACC1/2, and then malonyl-CoA and acetyl-CoA are used by FAS to generate palmitate (Fig 4A). To fix the efferocytosis defect in lipin-1 KO macrophages, we attempted to inhibit FA synthesis using c75, a synthetic FAS inhibitor (FIG 5B). However, inhibiting FAS did not reverse the efferocytosis defect in lipin-1 KO macrophages (FIG 5C). Interestingly, FAS inhibition impaired the efferocytic capacity of WT macrophages, indicating that basal production of fatty acids is required for efferocytosis (FIG 5C). Inhibition of ACC with Cpd9 showed similar trends in inhibiting efferocytosis in WT macrophages and a failure to restore efferocytic capacity in lipin-1 KO BMDMs (FIG 5C).

We previously demonstrated that loss of lipin-1 led to increased amounts of citrate/isocitrate in IL-4-stimulated BMDMs (*23*). Citrate can activate and fuel the FA synthesis pathway; therefore, we inhibited the CTP/citrate carrier to prevent citrate influx into the fatty acid biosynthesis pathway. Citrate carrier inhibition restored efferocytosis capacity in lipin-1 KO BMDMs without impairing WT macrophages (FIG. 5C). We believe that this restoration was due to the prevention of excessive fatty acid synthesis rather than a complete inhibition of lipid synthesis, which is necessary for membrane remodeling that is required during efferocytosis. To further demonstrate that impaired efferocytosis in lipin-1 KO macrophages is due to excess citrate export that can be utilized for FA synthesis, we treated WT macrophages with increasing amounts of exogenous citrate that have been previously shown to augment proinflammatory responses in LPS-stimulated monocytes (*53*) and increase lipid synthesis (*54*). Our data shows that exogenous citrate impairs the efferocytosis function of WT macrophages in a dose-dependent manner. (FIG 5D)

Excess-free fatty acids can be toxic to the cell, and when present in large amounts, they are directed toward the de novo sphingolipid pathway (*55*, *56*). This pathway leads to the production of ceramides, which impair the function of mitochondria, pro-resolving macrophages, and efferocytosis (*48*, *50*, *51*). Our results indicate that the excessive synthesis of fatty acids is directed toward the de novo sphingolipid pathway. To address this, we inhibited serine palmitoyltransferase (SPT), the enzyme responsible for the first step of ceramide biosynthesis. SPT catalyzes the reaction between palmitoyl-CoA and L-serine to produce ceramides. Inhibiting SPT with myriocin restored the efferocytic capacity of lipin-1 KO BMDMs (Fig 5E). Therefore, our data suggest that the buildup of ceramides due to increased fatty acid synthesis is responsible for the impairment of efferocytosis in lipin-1 KO BMDMs.

### Citrate carrier inhibition depletes neutral lipids without restoring mitochondria function

Beta-oxidation and oxidative metabolism have been attributed to promoting efferocytosis. In contrast, the contribution of the anabolic arm of lipid metabolism has been less studied (*21*, *23*). Excessive FA-induced ceramide production can impair macrophage efferocytosis function (*50*, *51*). Since several studies have shown that ceramides inhibit mitochondrial function and oxidative phosphorylation (OXPHOS) (*48*, *57*), we hypothesized that citrate carrier inhibition in macrophages promoted mitochondrial function. Consistent with previous cancer studies where inhibition of citrate carrier impaired mitochondrial respiration (*58*), we report that citrate carrier inhibition failed to restore the mitochondrial respiratory capacity in lipin-1 KO macrophages (Fig 6A). Since citrate carrier inhibition restored efferocytosis in lipin-1 KO macrophages without restoring mitochondrial respiration, this suggests that mitochondrial defect in lipin-1 KO macrophages is upstream of lipid synthesis.

**Fig. 6.**
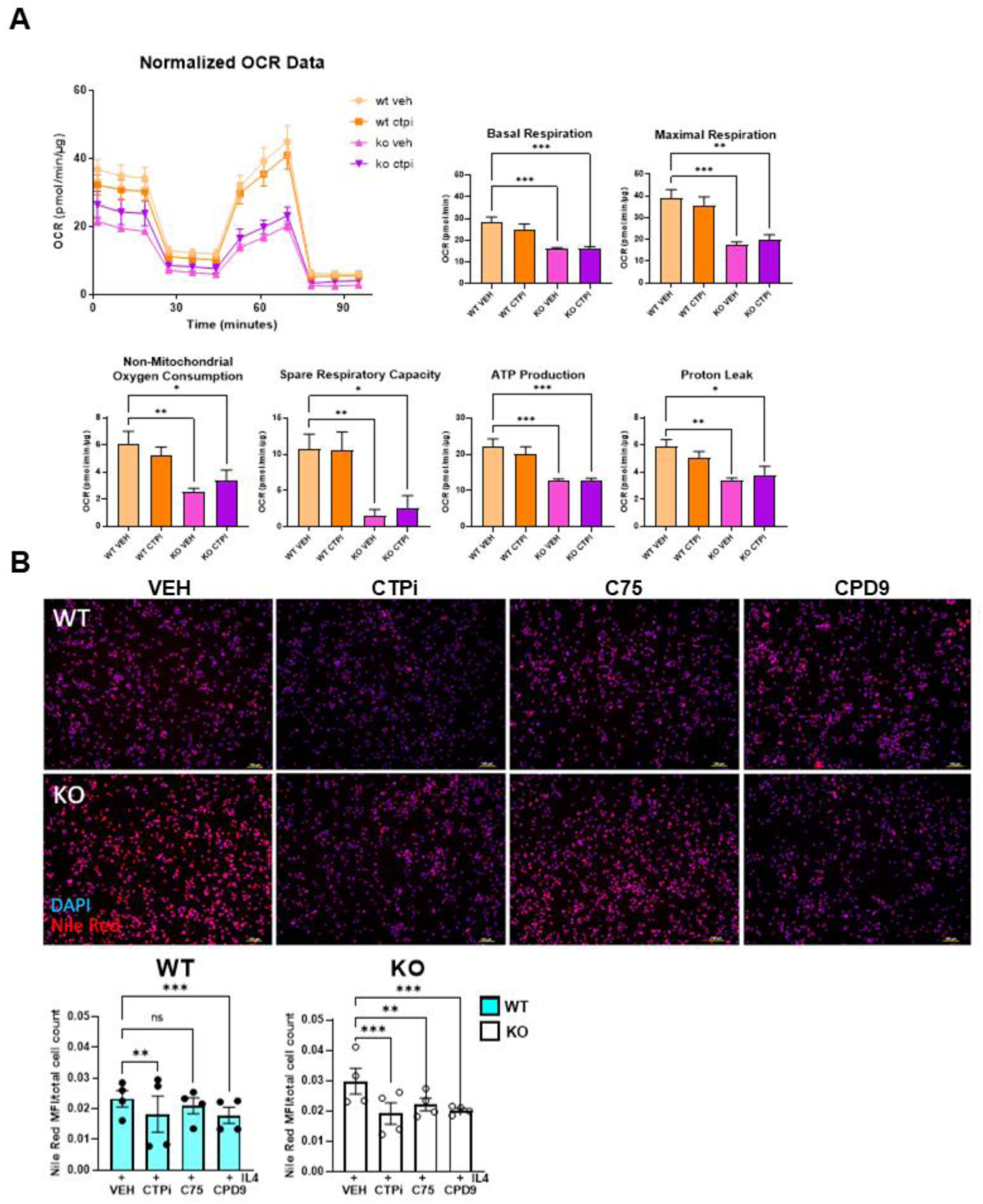
Inhibition of citrate carrier reduced neutral lipids levels without restoring mitochondria respiratory capacity. (**A**) BMDMs isolated from lipin-1mKO mice and littermate controls were treated with CTPi for 12 hours before cotreatment with CTPi and 40ng/ml IL-4 for 4 hours. Oxygen consumption rate (OCR), and mitochondrial function parameters were analyzed via Seahorse extracellular flux analyzer. Graphed data represent mean OCR with SEM (n=4) (**B**) BMDMs were treated with 10µM inhibitors (FAS: C75, ACC: Cpd9, CTP: CTPi) and DMSO Vehicle (VEH) for 12hrs and subsequently cotreated with 40ng/ml IL4 and 10µM of respective inhibitors for 4 hours before Nile red staining. Images were taken at 10x to generate representative images. The experiment was done twice, each time with 2 unique pairs of individual WT and FKO (full KO) BMDMs (n=4). At least 3 random images of each group were taken, quantified, and grouped to give individual dots. Bars represent the standard error of the mean (± SEM). Significance was determined by one-way and two-way ANOVA for Seahorse data and Nile red values, respectively. *= p≤0.05, **= p≤0.01, ***= p≤0.001.

Since inhibiting the cytoplasmic citrate influx should reduce FA biosynthesis and downstream lipids, we treated WT and lipin-1 KO macrophages with lipid synthesis inhibitors and stained them for neutral lipids. As expected, we observed a significant decrease in neutral lipids in lipin-1 KO macrophages with CTP, C75, and Cpd9 inhibition (Fig 6B). Even though WT macrophages start with reduced amounts of neutral lipids, similar trends were observed in WT macrophages except for C75, which only showed a decreasing trend in neutral lipid levels (Fig 6B). Though we hypothesized that there would be fewer lipids with FAS and ACC inhibition, we acknowledge the limitation of Nile red staining as it does not detect global lipids and cannot differentiate between intracellular and membrane-neutral lipids composition.

### Citrate carrier activity is responsible for defects in inflammation resolution

We have previously demonstrated that deletion of myeloid-associated lipin-1 delayed full skin wound closure and increased atherosclerotic plaque and necrotic core build-up (*32*, *38*). In addition, this study also demonstrated that loss of lipin-1 impairs inflammation resolution. Efficient efferocytosis is critical for the resolution of inflammation (*5*), and the inhibition of citrate carrier restored efferocytosis in macrophages deficient in lipin-1; thus, we hypothesized that inhibition of the citrate carrier in lipin-1^m^KO mice could restore inflammation resolution. Similar to the experiment in Fig 1A, we used the zymosan-induced peritonitis model and quantified neutrophils after 24 hours of treatment, as this time-point is where we observed the most significant difference in neutrophil numbers (Fig 1C). Following injection with zymosan, lipin-1^m^KO mice and littermate controls were injected with 50mg/kg of CTPI-2, a more potent and *in vivo*-suited second-generation CTP inhibitor at 12 and 18 hours before collection of cells by I.P lavage at 24 hours for flow cytometric analysis (Fig 7A). Consistent with the restoration of efferocytosis via citrate carrier inhibition, CTP-2 restored inflammation resolution in lipin-1^m^KO mice. Collectively, lipin-1 restrains FA synthesis to promote pro-resolving macrophage efferocytosis, which is required to resolve inflammation.

**Fig. 7.**
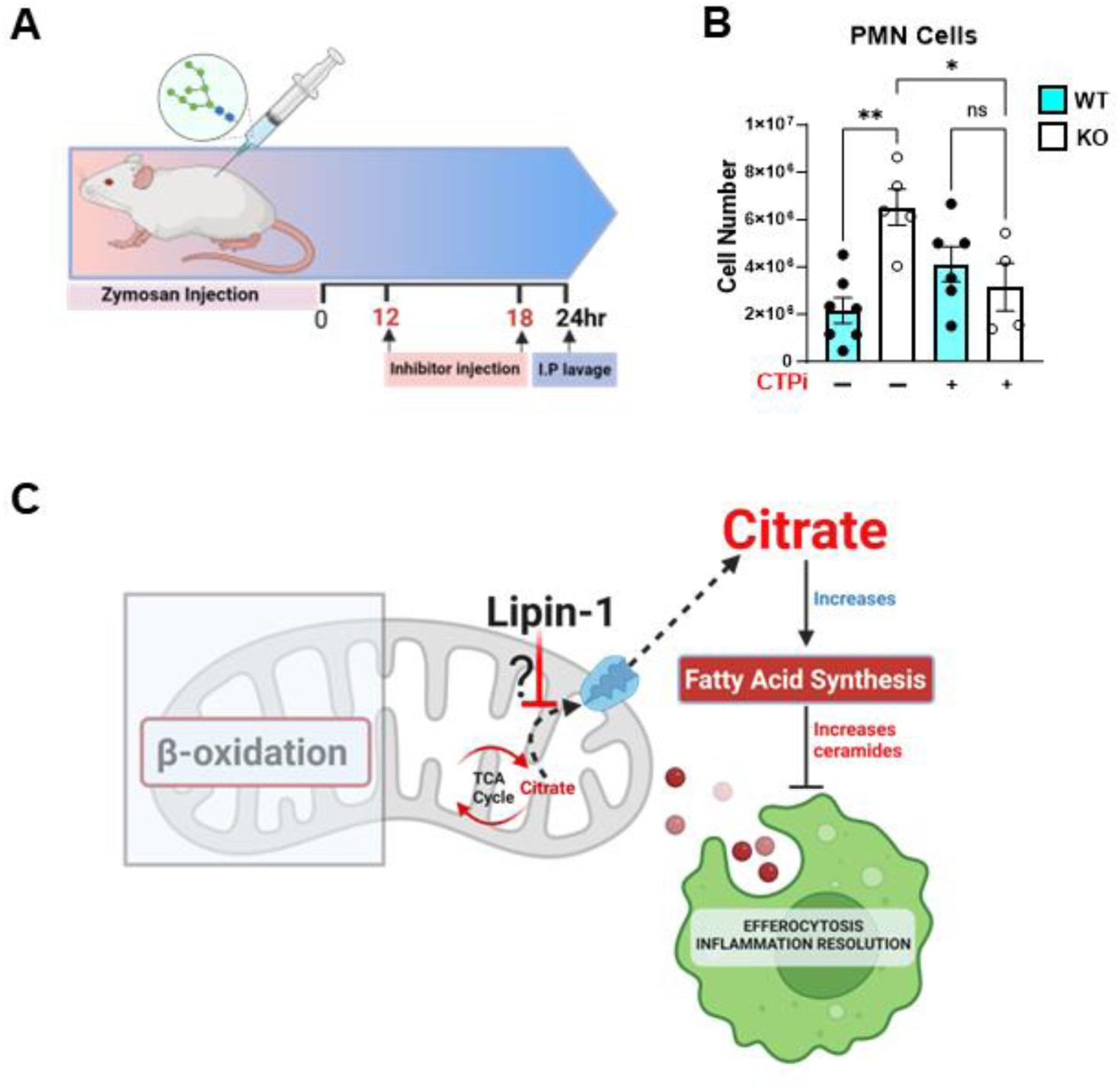
Inhibition of the citrate carrier restores inflammation resolution in lipin-1^m^ko mice. (**A**) Lipin-1^m^KO mice and littermate controls were injected with 50mg/kg CTP-2 or C75 at 12 hours and 18 hours post Zymosan injection. (**B**) Flow cytometry analysis of intraperitoneal lavage after acute (0.1mg/mouse, 24 hours) Zymosan injection. Illustration in (**A** and **C**) was created using BioRender.com. Bars represent the standard error of the mean (± SEM). Significance was determined by one-way ANOVA. *= p≤0.05, **= p≤0.01.

## Discussion

Unresolved inflammation promotes the progression of numerous disease conditions (*1–3*). Macrophages facilitate disease resolution via efferocytosis and by secretion of anti-inflammatory mediators to restore tissue homeostasis (*21*, *59*). Macrophage metabolism influences their activation state and function. Hence, macrophages require an appropriate metabolic state that supports pro-resolving function (*6*, *7*). Several aspects of lipid metabolism, including fatty acid oxidation and biosynthesis, contribute to macrophage polarization and function (*17*, *18*, *23*, *60*) (*18*). Lipin-1 plays a critical role in regulating lipid metabolism. We have previously demonstrated that lipin-1 promotes disease resolution in studies where the loss of myeloid-associated lipin-1 led to delayed wound healing and worsened atherosclerosis (*32*, *38*). In this study, we utilized the lipin-1^m^KO mice and the Lipin-1^m^EnzyKO mouse to investigate the contribution of lipin-1 to inflammation resolution and further understand how lipin-1 regulation of macrophage metabolism supports pro-resolving macrophage responses. We provide evidence that the non-enzymatic activity of lipin-1 restrains FA synthesis to promote efferocytosis and inflammation resolution.

After an acute inflammatory response, macrophages must polarize into a pro-resolving phenotype to dampen the ongoing inflammation (*5*). To this effect, pro-resolving macrophages alter their metabolism to support inflammation resolution processes, such as producing anti-inflammatory mediators and preventing inflammatory cell death of neighboring cells by efferocytosis (*3*, *6*, *7*). In this study, we show that myeloid-associated lipin-1 promotes inflammation resolution by facilitating the clearance of neutrophils in a Zymosan model of peritonitis. As macrophages must attain a metabolic profile that supports pro-resolving macrophage functions, we investigated the contribution of lipin-1 to the metabolic profile of macrophages during the resolution phase of inflammation. We report that loss of lipin-1 in peritoneal macrophages promotes a metabolic profile similar to a pro-inflammatory macrophage rather than a pro-resolving macrophage (*10*, *11*, *16*). We observed a population of macrophages that was more prevalent in lipin-1^m^KO mice with decreased key metabolic markers such as CD36, CPT1A, Cytc, and SDHA. The low levels of CPT1A, Cytc, and SDHA in peritoneal macrophage clusters unique to lipin-1^m^KO mice suggest an impairment in the TCA cycle and mitochondria homeostasis. CD36 is a scavenger receptor known to recognize ACs (*61*, *62*). CD36 and the platelet-activating factor receptor (PAFR) are proposed to promote apoptotic cell recognition and engulfment (*62*). Stimulation of macrophages with apoptotic cells was also shown to promote CD36 and PAFR interaction at the macrophage plasma membrane with the possible coupled function of forming lipid rafts that facilitate efferocytosis (*62*). Though Parks *et al*. showed that the resolution of 24-hour Zymosan-induced peritonitis is independent of CD36 activity, loss of CD36 leads to a delay in inflammation resolution typified by a 7-day post-injury increase in neutrophils, apoptotic cells, and macrophage numbers in a murine model of bleomycin-induced lung injury (*61*). We attribute the impairment in inflammation resolution in lipin-1^m^KO mice to defects in pro-resolving macrophage functions. This observation is consistent with other studies correlating defects in pro-resolving macrophage functions to the progression of chronic inflammatory diseases such as atherosclerosis and myocardial infarction (*21*, *40*, *59*). In addition, the altered metabolic profiles of lipin-1 deficient pMACs are consistent with our previous *in vitro* studies, where we reported a dysfunctional TCA cycle and impaired mitochondrial function in macrophages deficient in lipin-1 when stimulated with IL-4 (*23*). Altogether, our data suggests that lipin-1 supports proper pro-resolving macrophage metabolism to promote inflammation resolution.

We investigated the lipid profile in macrophages in response to pro-resolving stimuli. Results from the lipidomics assay position lipin-1 as a significant regulator of macrophage lipid metabolism because the loss of lipin-1 led to major changes in the lipidomic profile of lipin-1 KO macrophages. The IL-4-dependent increase in neutral lipids and ceramides observed in macrophages deficient in lipin-1 suggests altered lipid channeling (*55*, *56*). Upon saturation of lipid stores, fatty acids are channeled into the sphingolipid biosynthesis pathway to contribute to plasma membrane integrity and other cellular functions (*55*). However, owing to their bioactive and cytotoxic properties, ceramides are produced minimally as needed or utilized to make inert membrane sphingolipids (*49*). Ceramides impair efferocytosis, reduce the respiratory chain efficiency, inhibit M2 polarization, and are highly enriched in atherosclerotic plaques(*48*, *50*, *51*, *63*, *64*). Notably, de novo synthesis of ceramides impairs efferocytosis by altering actin polymerization via inhibition of Rac1 plasma membrane recruitment, which decreases membrane ruffle formation (*51*). Efferocytosis-induced Rac1 activation promotes the formation and stabilization of the phagocytic cup, which supports the uptake of multiple apoptotic cells (*59*). Together with our data where myriocin treatment restored efferocytosis in lipin-1 KO macrophages, we further provide evidence of the detrimental nature of ceramides on efferocytosis and inflammation resolution and the importance of lipin-1 in proper lipid channeling.

Upon the export of citrate into the cytoplasm, a series of enzyme-catalyzed reactions lead to the synthesis of fatty acids. Our data investigating the FA biosynthesis pathway suggests that the basal expression of enzymes involved in the FA biosynthesis pathway is sufficient to handle a high substrate influx since we do not observe significant differences in the abundance of CIC and FAS. As the notable increase in lipid species is in specific lipid families and not globally, we believe the differences in the lipidome profile in macrophages deficient in lipin-1 result from impaired lipid handling. Furthermore, we believe the increase in FA and other neutral lipids that we observe in lipin-1 KO macrophages is not due to persistent activation of de novo FA biosynthesis but possibly because of increased cytoplasmic citrate export and failure to restrain FA biosynthesis at key checkpoints, particularly at the rate-limiting step. This might explain why we see no difference in the protein levels of FAS. Though lipin-1 in non-myeloid cells is known to repress SREBPs, the master regulators of lipid metabolism via its transcriptional regulatory activity, citrate is a known allosteric activator of ACC, and we have previously reported an increase in citrate in macrophages deficient in lipin-1. In addition, while one would expect the inverse outcome, more buildup of lipids in WT macrophages, since WT macrophages retain the enzymatic activity of lipin-1 that can dephosphorylate phosphatidic acid to make DAG and TAG further downstream, there is the possibility that lipin-1 KO macrophages use monoacylglycerol acyltransferase and possibly, other novel enzymes to generate DAG. In addition, other studies have shown that loss of lipin-1 leads to increased lipid synthesis (*29–31*). We think lipin-1 preferentially regulates lipid metabolism to align with macrophage cellular function.

Lipid metabolism is self-regulating, and a negative correlation exists between lipid catabolism and anabolism (*65*). Efficient beta-oxidation has been shown to promote efferocytosis (*21*, *23*); thus, potentially inhibiting FA synthesis may restore efferocytosis. We demonstrated that CIC inhibition restored efferocytic capacity in lipin-1 KO macrophages and reduced the amount of neutral lipids. Additionally, CIC inhibition restored inflammation resolution in lipin-1^m^KO mice. CIC levels are high in Non-alcoholic steatohepatitis (NASH) livers (*66*). In a study that investigated the role of the citrate carrier in a model of Nonalcoholic fatty liver disease (NAFLD), it was shown that CIC inhibition reverted hepatic steatohepatitis, normalized glucose intolerance, and reduced inflammation with subsequent induction of an anti-inflammatory response (*66*). Inhibition of CIC also reduced liver mRNA and protein levels of ACC, SREBP, and FAS (*66*). Consistent with our findings, Tan *et al*. also showed that loss of CIC led to decreased liver fat, palmitate, monoglycerides, DAGs, and TAGs (*66*). Other studies have also reported the inflammatory role of the mitochondrial citrate carrier and its contribution to altered lipid biosynthesis (*12*, *53*, *67*). In macrophages, the CIC has been shown to be required for inflammatory cytokine induction of nitric oxide and prostaglandins (*12*). CIC-mediated cytokine, nitric oxide, and ROS production have also been implicated in the pathophysiology of COVID-19 (*68*). Thus, our data suggests that CIC inhibition reduces the build-up of lipids in lipin-1 KO macrophages to restore proper efferocytic function.

Much emphasis has been placed on the role of beta-oxidation in supporting pro-resolving macrophage function and efferocytosis. It is important to realize that beta-oxidation and FA biosynthesis are interconnected and that lipid metabolism is self-regulating since FA synthesis inhibits β-oxidation, and β-oxidation exhausts substrates required for FA synthesis (*65*). Consequently, studies that have reported impairment in efferocytosis due to pharmacological inhibition or genetic ablation of essential beta-oxidation proteins need to be open to the possibility that inhibition of beta-oxidation increases the acetyl-CoA pool that becomes available for lipid biosynthesis. This build-up in lipids, particularly ceramides, could then be responsible for inhibiting efferocytosis, which is usually attributed to beta-oxidation impairment. At the same time, in studies where beta-oxidation agonists have been used to promote efferocytosis, the underlying mechanism could result from decreased acetyl-CoA pool available for lipid biosynthesis due to increased beta-oxidation efficiency. Consistent with this notion, inhibiting the mitochondrial citrate carrier restored efferocytosis function in lipin-1 KO macrophages without restoring mitochondria function. In addition, citrate carrier inhibition in cancer cells decreases the respiratory chain complex-I activity and induces membrane potential destabilization, mitochondria fragmentation, and depletion by mitophagy (*58*, *69*). Thus, cell type differences could exist since citrate carrier inhibition does not impair WT macrophage mitochondrial function. Another possibility is that, in contrast to WT macrophages in our study, a more noticeable reduction would be observed in cancer cells’ mitochondria respiration since cancer cells already express a very high amount of the citrate carrier to support oxidative metabolism (*58*, *69*). The inability of ACC and C75 inhibition to restore efferocytosis supports the notion that a basal amount of lipid biosynthesis is required for efferocytosis(*19*, *62*). While our study suggests an inhibitory role of FA synthesis during inflammation resolution, FA synthesis is critical to mount an effective antimicrobial or pro-inflammatory response (*17*). Notably, the loss of ACC in macrophages impaired the ability to mount a proinflammatory response and phagocytic clearance of bacteria, which exacerbated the outcome of murine bacteria peritonitis (*70*).

While the findings in this study are sufficient to make inferences and draw conclusions, we understand that this study has limitations. Because of their low signal ratio, certain lipid species were undetectable or excluded in our lipidomics assay. We also understand that our in vivo studies were done in transgenic mice lacking full-length lipin-1 or the enzymatic domain of lipin-1 in myeloid cells. Thus, there might be other contributory aspects of other cells of myeloid origin. With this in mind, we demonstrate macrophage-specific differences throughout this study. While the efficacy of the mitochondrial citrate carrier has been demonstrated in cancer and NASH studies (*58*, *66*), our future direction will be to investigate the effect of the citrate carrier in the amelioration of atherosclerosis and other cardiometabolic diseases by employing pharmacological and genetic depletion/ablation experimental strategies.

We have shown that the role of lipin-1 in restraining FA and resolving inflammation is independent of its enzymatic activity. These findings align with our previous study, showing that lipin-1 promoted oxidative metabolism and efferocytosis independent of its enzymatic activity. (*23*). Aside from its enzymatic and transcriptional regulatory activity, lipin-1 can be associated with other proteins and on membranes and diverse organelles (*71*). Lipin-1 could also have a scaffolding or tethering function to facilitate lipid handling and anaplerotic function. This idea will also form the basis of our future investigations to delineate which non-enzymatic activity of lipin-1 is responsible for our current findings. In conclusion, the ability to restore efferocytosis and inflammation resolution independently of improved mitochondria function with CIC inhibition suggests that impaired mitochondria function in lipin-1 KO macrophages is upstream of altered lipid metabolism and possibly that the toxic downstream effect of FA biosynthesis is more crucial to the efferocytic process than efficient mitochondria function. Our findings strongly suggest that in response to pro-resolving stimuli, lipin-1 restrains FA synthesis, thus preventing the downstream production of ceramides that impair pro-resolving macrophage function and activities required for inflammation resolution (Fig 7C).

## Material and methods

### EXPERIMENTAL DESIGN

We used transgenic mouse models to investigate the role of macrophage-associated lipin-1 in resolving inflammation and its impact on lipid metabolism for promoting pro-resolving macrophage response. Bone marrow-derived macrophages were generated from age-matched lipin-1mEnzyKO, lipin-1mKO, and their littermate controls. In vitro experiments used primary cells from multiple mice, with technical replicates as needed. In Zymosan-induced peritonitis in vivo experiments, investigators were blinded to mouse genotypes, and mice were randomly assigned to groups. Each experimental condition included at least three KO and WT mice. Health deterioration was monitored during the mild and self-limiting zymosan-induced peritonitis experiments. All animal studies were approved (ACUC# P22-014).

### MICE

All mice studies were approved by LSU Health Shreveport institutional animal care and use committee (ACUC# P22-014). Mice were housed and cared for according to the National Institute of Health guidelines for the care and use of laboratory animals. Mice lacking lipin-1 enzymatic activity from myeloid cells (lipin-1^mEnzy^KO) were generated as previously reported (*34*). Briefly, mice with exons 3 and 4 of the *Lpin1* gene flanked by LoxP sites (genetic background: C57BL/6J and SV129; generously provided by B.N.F. and R. Chrast) were crossed with C57BL/6J LysM-Cre transgenic mice purchased from The Jackson Laboratory (Bar Harbor, ME). Mice entirely lacking lipin-1 from myeloid cells (lipin-1^m^KO) were generated by crossing mice with exon 7 of the *Lpin1* gene flanked by LoxP sites (genetic background: C57BL/6J and SV129; generously provided by B.N.F (*72*) with C57BL/6J LysM-Cre transgenic mice purchased from The Jackson Laboratory. Age-matched lipin-1 flox/flox littermate mice were used as controls. Mice were genotyped to detect Cre insertion using primer combinations based on Jackson lab protocols and the following primers (AAGGAGGGACTTGGAGGATG-common forward, GTCACTCACTGCTCCCCTGT-wild type reverse, ACCGGTAATGCAGGCAAAT-mutant reverse)

### BONE MARROW-DERIVED MACROPHAGE GENERATION

Mice femurs were harvested and flushed with unsupplemented DMEM (Dulbecco’s Modified Eagle Medium, Gibco, 11965-092) to isolate bone marrow. Bone marrow cells were purified through a series of centrifugations. Red blood cells were removed by lysing bone marrow cells with ammonium chloride-potassium carbonate lysis (1X ACK lysis buffer), followed by filtration. Purified bone marrow cells were then resuspended and incubated for 7 days in non-treated T175 tissue culture flasks at 37°C and 5% CO_2_ in BMDM differentiation media composed of KnockOut^TM^ DMEM (Gibco, 10829-018), 30% L-cell conditioned medium (generated from L929 cells; ATCC CCL-1), 1mM sodium pyruvate (CORNING, 25-000-CI), 2 mM GlutaMAX^TM^ (Gibco, 35050-061), 100 U/ml penicillin-streptomycin (Gibco, 15140-122) 0.2% sodium bicarbonate (Gibco, 25080-094) and 20% FBS (ATLAS biologicals, EF-0500-A). To harvest macrophages, after cells reach 80% confluency, BMDM differentiating media was removed, cells were washed with DPBS (Dulbecco’s Phosphate-buffered saline without calcium and magnesium; 14190-144), then subsequently incubated in 11mM pH 7.6 EDTA (Fisher Scientific, BP120-1) in DPBS (Corning, Cat #21-031-CV) to detach cells. BMDMs were then placed in D10 media containing DMEM (Gibco, 11965-092), 10% fetal bovine serum (ATLAS biologicals, EF-0500-A), 2 mM GlutaMAX^TM^ (Gibco, 35050-061), 100 U/ml penicillin-streptomycin (Gibco, 15140-122) and 1mM sodium pyruvate (CORNING, 25-000-CI)

### L-CELL CONDITIONED MEDIUM

The murine fibroblast cell line L929 (ATCC CCL-1) was grown in RPMI Medium 1640 (Gibco, 11875-093) supplemented with 10% FBS (ATLAS biologicals, EF-0500-A), 2 mM GlutaMAX^TM^ (Gibco, 35050-061), 1mM sodium pyruvate (CORNING, 25-000-CI), and 100 U/ml penicillin-streptomycin (Gibco, 15140-122). Briefly, 3.75x10^5^ L929 cells were seeded in a T225 tissue culture flask with 75 mL supplemented RPMI media. Flasks were incubated for 12-14 days at 37°C until cells were 100% confluent and formed a cobblestone appearance. The medium was collected, cleared of cell debris by centrifugation at 1600RPM, filtered (0.2 µm), stored at 4°C for 24 hours, 0°C FOR 24 hours, and finally at -80°C until use.

### APOPTOTIC CELL GENERATION

Jurkat cells (ATCCVTIB-152™) cultured in RPMI Medium 1640 (Gibco, 11875-093) supplemented with 10% FBS (ATLAS biologicals, EF-0500-A), 2 mM GlutaMAX^TM^ (Gibco, 35050-061), 10mM HEPES buffer solution (Gibco, 15630-080), and 100 U/ml penicillin-streptomycin (Gibco, 15140-122) were centrifuged to remove culture media, resuspended in PBS. 10mls of Jurkat cells (1-1.5 x 10^6^ cells/ml) were seeded in 100mm x 15mm petri dishes (Fisher Scientific, FB0875712) exposed to UV radiation for 5 mins in the UV Clave™ Ultraviolet Chamber (Benchmark Scientific). PBS containing Jurkat cells was collected, and 6ml of 11mM EDTA (pH 7.6) was added to the Petri dishes for 10 minutes to detach the adherent cells and added to the initial pool of UV-radiated cells. Cells were centrifuged at 1600RPM for 5 mins, resuspended in supplemented (O% FBS) DMEM, and incubated for 2 hours at 37°C before downstream experimental use.

### EFFEROCYTOSIS ASSAY

1.25 x 10^5^ BMDMs from lipin-1^m^KO mice and littermate controls were seeded on 12mm coverslips (Fisher Scientific, 12541002), allowed to sit for 3 hours, and treated with recombinant mouse IL-4 (Leinco technologies, Inc. I-207) at a 40ng/ml concentration for 6 hours. Following IL-4 stimulation, macrophages were challenged with CFSE (5-[and 6]-Carboxyfluorescein diacetate succinimidyl ester Kit, BioLegend,423801)-labeled apoptotic cells at a 4:1 (AC/Macrophage) ratio for 45 mins. Unbound ACs were washed off with PBS, and cells on coverslips were fixed with 10% Neutral Buffered formalin (VWR, 16004-128) for 1hr. Fixing was stopped through a series of PBS washes. Cells on coverslips were stained with DAPI (1:36400, Invitrogen, D3571) for 15 mins, mounted on slides, and imaged on the BZ-X810 Keyence microscope (Keyence Corporation of America, Itasca, USA) at 20x. The number of macrophages was counted with the Keyence analyzer software, and the number of engulfed ACs was counted with the image J cell counter (Rasband, W.S., ImageJ, U. S. National Institutes of Health, Bethesda, Maryland, USA, https://imagej.nih.gov/ij/, 1997-2018). For inhibitor studies, cells were pretreated for 12 hours with 10µm CTPi (Calbiochem, 475877-5MG) or 10µm C75 (Sigma-Aldrich C5490-5MG) or 10µm CPD9 (Calbiochem^®^, 5343350001) or for 2 hours with 10µm Myriocin (MedChemExpress, HY-N6798) before 6 hours cotreatment with 40ng/ml IL-4 and respective inhibitor. In the citrate complementation experiments, the experimental setup was modified from (*53*). Cells were pretreated with increasing concentrations (1mM, 3mM, 6mM) of sodium citrate tribasic (SCB) dihydrate (Sigma-Aldrich 71402-250G) for 2 hours before 6 hours cotreatment with 40ng/ml IL-4 and SCB at respective concentrations.

### NILE RED ASSAY

1.75 x 10^5^ BMDMs from lipin-1^m^KO mice and littermate controls were seeded on 12mm coverslips (Fisher Scientific, 12541002) or in tissue culture treated 24 well plates, allowed to sit for 3 hours and treated with recombinant mouse IL-4 (Leinco technologies, Inc. I-207) for 4 hr. For inhibitor studies, cells were pretreated as aforementioned. After IL-4 stimulation, cells were fixed with 10% Neutral Buffered formalin (VWR, 16004-128) for 1hr. Following PBS washes, cells were stained with 0.5ug/ml Nile red (Invitrogen N1142) for 10 minutes, washed, and subsequently stained with DAPI (1:36400 dilution, Invitrogen, D3571) for 15 mins. Cells were imaged on the BZ-X810 Keyence microscope (Keyence Corporation of America, Itasca, USA). Macrophage numbers were counted using the BZ-X800_Analyzer (Keyence Corporation of America, Itasca, USA), and Nile red fluorescent intensity was quantified using cellSens Dimension 1.16 (OLYMPUS, Tokyo, Japan)

### IN VIVO PERITONITIS

lipin-1^m^KO mice, lipin-1^m^EnzyKO, and littermate controls were injected with 0.5cc (0.1mg/mouse) of 0.2mg/ml Zymosan (ZYMOSAN A from *Saccharomyces cerevisiae,* Sigma-Aldrich, Z4250) solution. After 4, 12, 24, and 48 hours, 5mls of sterile, cold FACS wash buffer (1% BSA and 0.1% sodium azide in PBS) was injected intraperitoneally into the mice and collected back as the peritoneal lavage. Cells in the peritoneal lavage were counted and prepared for flow cytometry staining. In detail, 5 x 10^5^ peritoneal cells were blocked with CD16/CD32 (1:200) for 25 mins at 4°C. Cells were treated with LIVE/DEAD Aqua (Invitrogen L34957) according to the manufacturer’s instructions, followed by incubation with AF700-conjugated anti-CD45.2 (1:2000) (109821, clone 104; BioLegend), PECy7-conjugated anti-CD11b (1:4000) (25-0112-81, clone M1/70; eBioscience), PECy5-conjugated anti-F4/80 (1:400) (15-4801-80, clone BM8; Invitrogen), and FITC-conjugated anti-Ly6G (1:800) (551460, clone 1A8; BD Biosciences) for 30 min in the dark at 4°C. Excess antibodies were washed off from cells with FACS wash and centrifugation at 1600RPM for 5 mins. Cells were resuspended in 500µl wash, and the immune cell population was quantified using a NovoCyte Quanteon 4025 flow cytometer. Appropriate Fluorescence Minus One Controls (FMOCs) were used to identify positive populations, and Compensation controls (Comp Bead, Invitrogen; 01-2222-42) were included in the experimental setup to exclude spectral overlap. For inhibitor studies, mice were injected with 50mg/kg of CTPI-2 (MedchemExpress CAS:68003-38-3) at 12 hours and 18 hours posts Zymosan injection, and intraperitoneal lavage was collected at 24 hours and analyzed for immune cell population and distribution via flow cytometry. Data analysis was performed using NovoExpress (ACEA Biosciences).

### SPECTRAL FLOW CYTOMETRIC METABOLIC AND INFLAMMATION PROFILING

After Zymosan injection, peritoneal lavage was collected after 6 days, as described above. If required, the cells were treated with ACK lysis buffer to lyse the red blood cells, washed, and counted. The 27-color/29-parameter panel to look at the metabolic and inflammatory markers was adapted from (*41*). Antibodies used for spectral flow cytometry are listed in Table 1. Abcam lightning-link kits were used to conjugate the intracellular targets. For each experiment, 2x10^6^ cells per sample were seeded in the 96-well V-bottom plate (Corning, 3894). Cells were washed with DPBS (Corning, Cat #21-031-CV) and first stained with Zombie Ultraviolet fixable viability dye (1:1000; BioLegend, 423107) for 30 min at room temperature to exclude the dead cells. Fc receptors were then blocked by incubating cells in TruStain Fcx plus Fc block (1:100; BioLegend, 156604) in FACS buffer (DPBS containing 2% FBS and 2mM EDTA) at 4°C for 10 min. The surface staining was carried out in FACS buffer for 30 mins at 4°C using anti-mouse antibodies against CD11c, CD206, CX3CR1, CD36, and CD19. Following surface staining, cells were fixed with eBioscience Foxp3 fixation/permeabilization staining kit (Invitrogen, #00-5523-00) for 30 mins at 4°C, permeabilized in 1X permeabilization buffer overnight, and Fc-blocked for 15 mins at 4°C. Intracellular staining was done in 1x permeabilization buffer for 1hr at 4°C using anti-mouse antibodies against Glut1, PKM, SDHA, CPT1A, ACC1, Cytc and G6PD. After intracellular staining, cells were washed twice with permeabilization buffer and once with FACS buffer. The remaining surface targets were stained in FACS buffer for 30 mins at 4°C (Surface stain 2, Table S1). Brilliant stain buffer plus (BD Biosciences, 566385) and monocyte blocker (BioLegend, 426103) were added to the staining buffer whenever required. Single-color (using both cells and ultracomp beads), unstained, and fluorescence minus one control (FMOCs) were prepared for each staining experiment. Finally, cells were suspended in 200 μl of FACS buffer and acquired on Bigfoot 5-laser Spectral Cell Sorter (Invitrogen). Acquired samples were unmixed using Bigfoot Sasquatch software and analyzed with FlowJo software v10.9 (Tree Star Inc.).

High-dimensional analysis was done using the OMIQ software from Dotmatics (www.omiq.ai, www.dotmatics.com). Initial gating was done using FlowJo; this involved selecting leukocytes, removing doublets, and removing dead cells. The cell population of interest was identified and gated as CD45+ CD11b+ F4/80+ (CD3- CD19- Ly6G-). This gated cell population was downsampled to 90,000 cells per sample. After downsampling, FCS files were exported and uploaded to the OMIQ platform. First, the parameters were scaled (scaling type = Arcsinh, Cofactor = 400, Min= -800, Max = 100000). UMAP was run using the default settings (Euclidean distance function, nearest neighbors: 15, and minimum distance: 0.4). It was followed by phenograph clustering using the Euclidean distance function and K = 100. OMIQ was also used to calculate the median fluorescent intensity. Graphs were made in Prism 10, v10.2.0 (GraphPad Software Inc.).

### LIPIDOMICS ASSAY

1 x 10^6^ BMDMs from lipin-1^m^KO mice and littermate controls were seeded in 6 well tissue culture-treated plates. Three technical replicates were included per individual experiment. Macrophages were incubated for 3 hours and subsequently stimulated with 40ng/ml IL-4 for 4 hours. After IL-4 stimulation, culture media was aspirated, and each well was washed with 1ml of room temperature 0.9% sodium chloride solution (B. Braun Medical Inc, R5201-01). Saline solution was aspirated, and 400µl of ice-cold LC-MS grade methanol (Infinity Lab Ultrapure LC-MS Methanol, Agilent technologies; 5191-4497) was added to each well and allowed to sit for 5 mins, after which 400µl of LC-MS grade water (Pierce^TM^ Water, LC-MS Grade, Thermo scientific; 51140) was added. Cells were scraped and transferred to a labeled polypropylene microfuge tube (Agilent Technologies, tube:5191-8150, caps:5191-8151). Samples were stored at -80°C before shipment to the Metabolomics core at the Rocky Mountain National Lab.

400µl of ice-cold LC-grade chloroform (Fisher ChemicalTM; C607-4) was added to each sample. Samples were shaken for 30 mins at 4°C and centrifuged at 16000g for 20 mins. 400 µl of the bottom (organic) layer was collected and dried in a Savant SpeedVac SPD130 (Thermo Scientific). Lipids were resuspended in 1 ml of 5 µg/ml butylated hydroxytoluene in 6:1 isopropanol:methanol.(Isopropanol, Fisher Chemical^TM^; A461-4; Methanol, Fisher Chemical^TM^; A456-4; butylated hydroxytoluene, MP Biomedicals™, 0210116280).

Bulk lipids were analyzed as previously described (*73*). Samples were separated using a Shimadzu Nexera LC-20ADXR HPLC and a Waters XBridge® Amide column (3.5 µm, 3 mm X 100 mm). Lipids were separated by headgroup with a 12-minute binary gradient from 100% 5 mM ammonium acetate, 5% water in acetonitrile pH_apparent_ 8.4 to 95% 5 mM ammonium acetate, 50% water in acetonitrile pH_apparent_ 8.0 (Water, Fisher Chemical^TM^, W64; Acetonitrile, Fisher Chemical^TM^, A996-4; Ammonium Acetate, Fisher Chemical^TM^, A11450). Lipids were detected using a Sciex 6500+ QTRAP® mass spectrometer with polarity flipping and scheduled MRMs.

All signals were integrated using MultiQuant® Software 3.0.3. Signals with greater than 50% missing values or an intensity of less than 3000 units were discarded, and the remaining missing values were replaced with the group average of 3000 units. All signals with a QC coefficient of variance greater than 40 % were discarded. A total of 731 individual lipid species remained in the final dataset. Data was the total sum normalized before analysis. Single and multivariate analysis was performed in MarkerView® Software 1.3.1.

### MITOCHONDRIAL BIOENERGETICS

1 x 10^5^ BMDMs were seeded in Agilent Seahorse XF24 Cell Culture Microplate (Agilent Technologies, 100777-004) and treated as required by experiments. After treatment, culture media was aspirated and replaced with warm SeaHorse assay media containing Agilent Seahorse XF DMEM Medium, pH 7.4 (Agilent Technologies 103575-100), 1mM pyruvate (Agilent, 103578-100), 2mM GlutaMAX^TM^ (Gibco, 35050-061), 10mM glucose solution (Agilent, 103577-100). Cells were incubated in a no CO_2_ incubator for 45 mins, and the Oxygen Consumption Rate (OCR) was measured by Agilent Seahorse XFe24 Analyzer (Agilent Technologies, CA, USA). Concentrations of the Mitochondrial stress assay inhibitors are as follows: 1µM oligomycin (Sigma-Aldrich, 75351-5MG), 2µM FCCP (Sigma-Aldrich, SML2959-1ML), 1µM Rotenone (Sigma-Aldrich, R8875-1G) and Antimycin (Sigma-Aldrich, A8674-25MG).

### WESTERN BLOTTING

Cells were lysed with 1X NUPAGE lysis buffer containing 1X Halt^TM^ protease inhibitor (Thermo Scientific, 18612793+9), 1X phosphatase inhibitor cocktail 2 (Sigma-Aldrich, P5726-5ML), 1X phosphatase inhibitor 3 cocktail (Sigma-Aldrich, P0044-5ML) and 100 mM dithiothreitol (VWR, 0281-25G). Peirce^TM^ 660 nm Protein Assay (Thermo Scientific, 22660) was used to determine the protein concentration of sonicated and heat-denatured protein samples. 10/20µg of protein sample was loaded in the wells of precasted 4-12% polyacrylamide NuPAGE Novex gel (Invitrogen, NP0322BOX) submerged in 1X MOPs running buffer. Protein bands were separated for 48 mins at 200 volts and 400 mAMPs. Subsequently, protein bands within the gel were transferred on an Immobilon-FL transfer membrane (Millipore IPFL00010) for 45 mins at 20 volts and 400 mAMPs. Membranes were blocked with PBS intercept buffer (Licor, 927-70001) for 1hr at room temperature. Membranes were incubated overnight with appropriate primary antibodies. Following primary antibody incubation, membranes were washed trice with 1x TBST (tris buffered saline with tween 20) before incubation with secondary antibody in 1x TBST containing 5% Non-Fat Dry Milk Omniblok (Americanbio, AB10109-01000) and 0.01% SDS (Thermo Scientific, 28365). Membranes were washed trice with 1x TBST before 1 min incubation with ImmunoCruz Western blotting luminol reagent (Santa Cruz, sc-2048). The Amersham Imager 680 (GE Healthcare Bio-Sciences) was used to capture protein bands, and the ImageQuant™ TL 8.1 (Cytiva) was used to quantify protein bands. For histone Immunoblot 5 - 15 µg of protein sample was loaded in the wells (equally) in 15% SDS-PAGE tris-glycine gels and run for 30 min at 200 V. Proteins were then transferred to nitrocellulose membranes (GVS NitroBind™. EP2HY00010) for 1hr at 90 V. Blocking was performed using Odyssey Intercept blocking buffer (LI-COR 92760001) for 1 h at room temperature. Primary antibody incubations were also performed using the Odyssey Intercept blocking buffer. After washing, membranes were incubated with a 1:15,000 dilution of LI-COR secondary antibodies. Membranes were washed four times in 1× Tris-buffered saline with Tween 20 (TBST). Membranes were stored in 1× TBS, imaged using an Odyssey infrared imaging system, and quantified using Bio-Rad Image Lab.

### WESTERN BLOT ANTIBODIES

Primary antibodies used for western blotting were Phospho-ACC (D7D11, CST: Cell Signalling and Technology), ACC (3662S, CST), P-AMPKalpha (T172, CST), AMPKalpha (2532S, CST), Lipin-1 (D2W9G, CST), CIC/CPT/SLC25AI (15235-I-AP Proteintech), AceCS1 (D19C6, CST) Fatty Acid Synthase (C20G5, CST), GAPDH (14C10, CST), Actin (A2066-2ML, Sigma), Pan H3 (Sigma-Aldrich #05-928), Pan H4 (Sigma-Aldrich #05-858), Acetyl H3 (EMD Millipore CORP #06-599), Acetyl H4 (Sigma-Aldrich #06-866). The working stock of all primary antibodies was made at 1:1000 dilution, except GAPDH and ACTIN, which were made at 1:5000 and 1:2000 dilution, respectively. Secondary antibodies used were Goat anti-rabbit IgG (111-035-003 Jackson ImmunoResearch, 1:2000) and Goat anti-mouse (610-1319 Rockland, 1:2000)

### STATISTICAL ANALYSIS

GraphPad Prism 9 (GraphPad Software, LLC) was used for statistical analyses. We carried out a test for normal distribution using the D’Agostino & Pearson test and Shapiro-Wilk test. For comparison between two data sets that passed the normality test, we used Student’s t-test analysis. A one-way ANOVA was used to analyze grouped data sets, while a two-way ANOVA was used to analyze grouped data sets with >2 technical replicates averaged as single values. Data were presented as standard error of the mean (± SEM), and statistical significance was assigned at *P < 0.05. Details of the statistical analysis of individual experiments are included in the figure legend.

## Acknowledgments

We acknowledge the services offered by the LSU Health Shreveport Research Core facility and the Center of Applied Immunology and Pathological Processes Immunophenotyping Core.

## Funding

This work was supported by the following grants:

National Institutes of Health R01HL163106 (MDW)

National Institutes of Health P20GM134974 (MDW, RSS, SB)

National Institutes of Health R01HL131844 (MDW)

National Institutes of Health R00HL145131 (AYJ)

National Institutes of Health R01HL167758 (AYJ)

National Institutes of Health R01HL119225 (BNF)

Ike Muslow Predoctoral Fellowship Intramural Award (TTB)

Intramural Research Program of the NIH, National Institute of Allergy and Infectious Diseases. (CMB)

## Author contributions

Conceptualization: TTB, RMS, OOI, MDW

Data curation: TTB, RMS, OOI, EHN, SB, BS, MDW

Methodology: TTB, RMS, OOI, EHN, SB, CD, BS, EB, RSS, AYJ, AYJ, MDW

Investigation: TTB, RMS, OOI, EHN, SB, BS, EB, MDW

Formal Analysis: TTB, RMS, OOI, EHN, BS, MDW

Visualization: TTB, OOI, SB, BS, MDW

Validation: TTB, SB, BS, MDW

Supervision: MDW

Resources: CMB, RSS, BNF, AYJ, MDW

Writing—original draft: TTB, RMS, MDW

Writing—review & editing: TTB, RMS, EHN, CD, BS, CMB, BNF, RSS, AYJ, MDW Project administration: MDW

## Data and materials availability

All data needed to deduce conclusions from this research article are included in the main manuscript and supplemental materials. Graphical abstracts and illustrations were created with BioRender software (https://app.biorender.com).

