## Supplemental Figures and Table for "Lipin-1 restrains macrophage lipid synthesis to promote inflammation resolution"

Temitayo T. Bamgbose *et al.*

**This PDF file includes:**

Figs. S1 to S3
Table S1

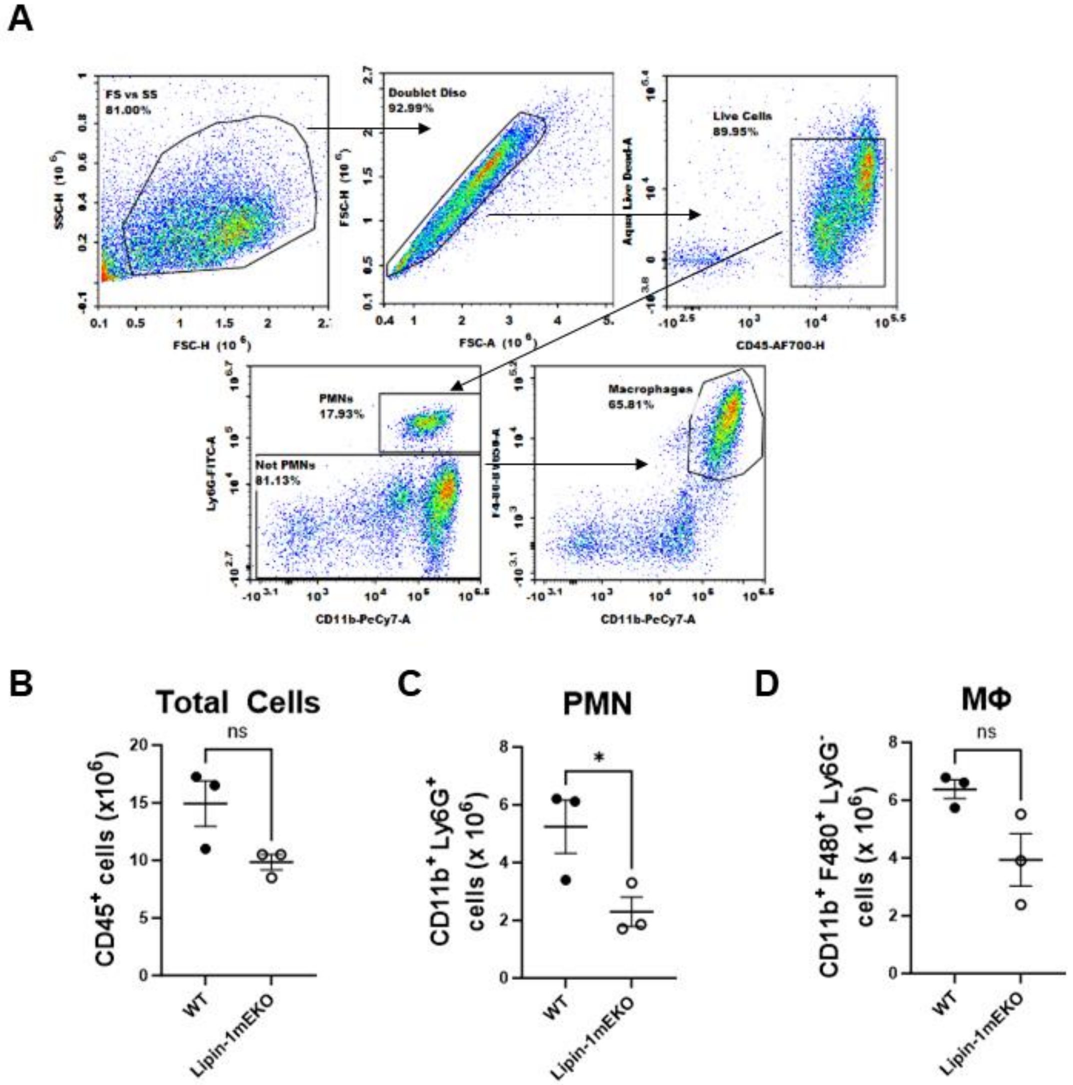


**Fig. S1.**

Mice were subjected to zymosan challenge (0.1mg/mouse). PMNs and macrophages were quantified from the peritoneal cavity by flow cytometry.

**A**. Gating strategy to remove doublets, dead cells and identify PMNs and Macrophages.

**B**. Total number of cells isolated from the peritoneal cavity.

**C**. Total number of PMNs isolated from the peritoneal cavity.

**D**. Total number of Macrophages isolated from the peritoneal cavity. A minimum of three mice per group per time point were analyzed. N $\geq$3 mice per group per time point. Values are means ± SEM. Unpaired Two-tailed T-tests were performed between groups at each time point. *=p$\leq$0.05

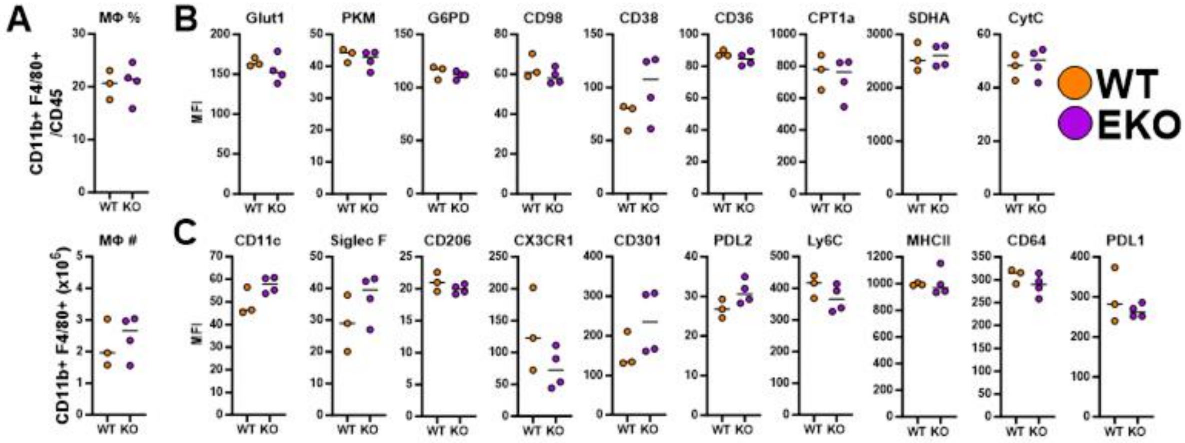


**Fig. S2.**

Mice were challenged with 0.1mg zymosan, and six days later, peritoneal cells were isolated by lavage. Isolated cells were stained for flow cytometric analysis.

**A**. The number and ratio of CD11b+ F4/80+ lin- cells in the peritoneal cavity.

**B-C**. Median fluorescent Intensity of metabolic and inflammatory markers of CD11b+ F4/80+ lin- cells. N=3 mice per group. Dots represent individual mice; lines are means. Significance was determined by unpaired Two-tailed T-tests. *=$p \leq 0.05$

Abbreviations: **Glut 1**: glucose transporter, **PKM**: Pyruvate Kinase M2, **G6PD**: Glucose-6-phosphate dehydrogenase, **ACC1**: Acetyl CoA carboxylase 1, **CPT1A**: mitochondria fatty acid importer, **SDHA**: Succinate dehydrogenase, **CytC**: Cytochrome C, **PDL**: Programmed Death-Ligand

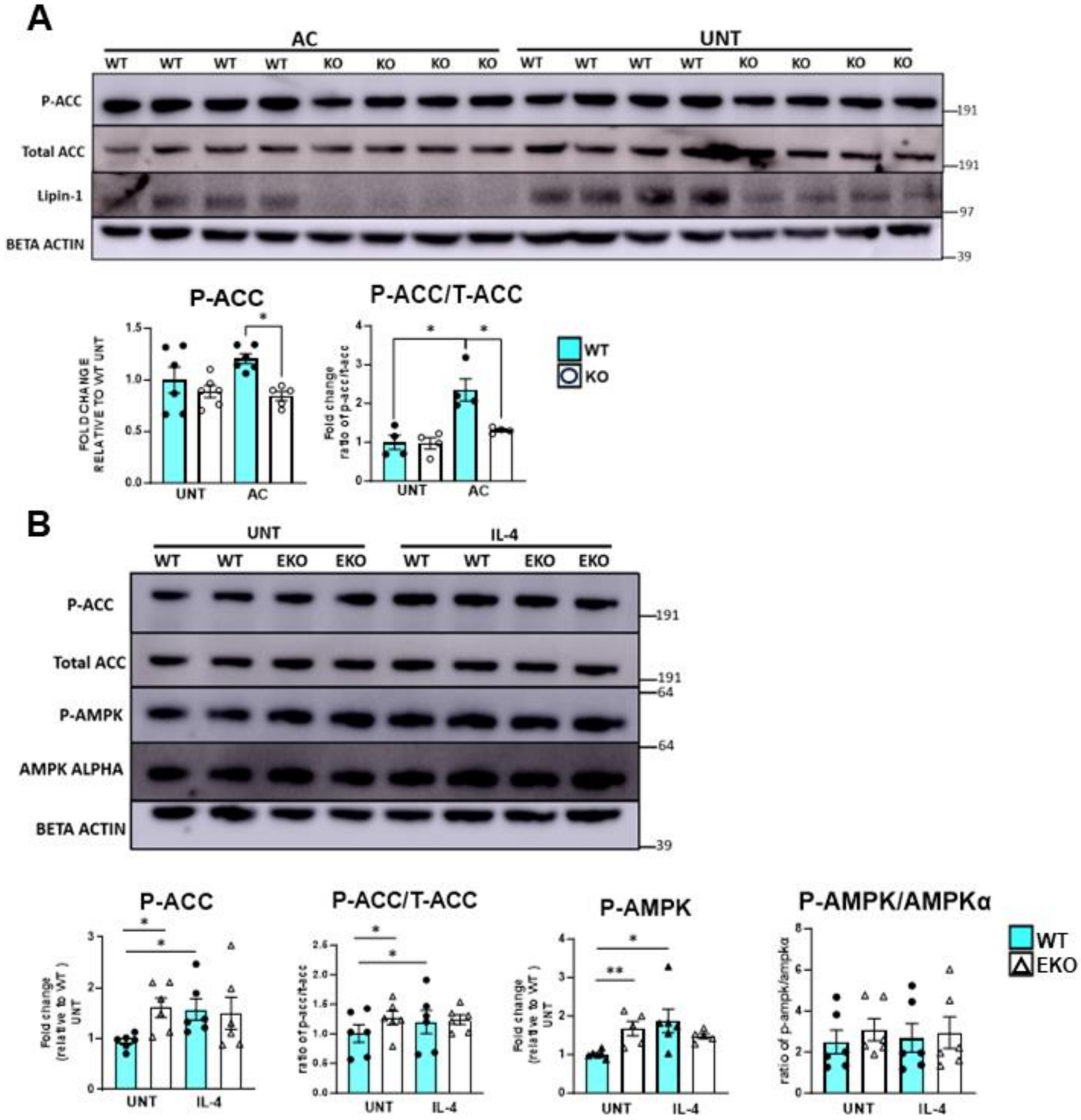


**Fig. S3.**

**A**. Macrophages were stimulated with AC at a 4:1 (AC/Macrophage) ratio for 45 mins, washed off and allowed to incubate in culture media for 3hrs and 15 mins.

**B**. Protein was isolated from IL-4-stimulated (40ng/ml, 4hrs) whole cells, and protein abundance of fatty acid biosynthesis enzymes was quantified by Western blot analysis (n>3). Bars represent the standard error of the mean (± SEM). Significance was determined by two-tailed student's t-test. *P < 0.05, **P < 0.01.

**Table S1. Spectral Flow Cytometric antibody panel**

| Target | Fluorophore | Company | Clone | Dilution | Identifier |
|---|---|---|---|---|---|
| CD11c | BUV496 | BD Biosciences | N418 | 1:400 | Cat: 750450<br>RRID: AB_2874611 |
| CD206 | BV785 | BioLegend | C068C2 | 1:400 | Cat: 141729<br>RRID: AB_2565823 |
| CX3CR1 | BV605 | BioLegend | SA011F11 | 1:100 | Cat: 149027<br>RRID: AB_2565937 |
| CD36 | AF700 | Thermofisher | HM36 | 1:100 | Cat: 56-0362-82<br>RRID: AB_2811887 |
| CD19 | BV510 | BioLegend | 6D5 | 1:100 | Cat: 115545<br>RRID: AB_2562136 |
| Intracellular stain | | | | | |
| Glut1 | LL-DL405 | Abcam | EPR3915 | 1:200 | Cat: ab252403 |
| PKM | PE | Abcam | EPR10138(B) | 1:1000 | Cat: ab210448 |
| SDHA | LL-AF647 | Abcam | EPR9043 | 1:1000 | Cat: ab240098 |
| CPT1A | LL-PE/Cy5 | Abcam | EPR21843-71-2F | 1:200 | Cat: ab235841 |
| ACC1 | LL-AF488 | Abcam | EPR23235-147 | 1:200 | Cat: ab272704 |
| Cytc | LL-PE/Cy7 | Abcam | 7H8.2C12 | 1:1000 | Cat: ab237966 |
| G6PD | LL-APC/Cy7 | Abcam | EPR20668 | 1:1000 | Cat: ab231828 |
| Surface stain 2 | | | | | |
| CD45 | APC/Fire810 | BioLegend | 30-F11 | 1:2000 | Cat: 103174<br>RRID: AB_2860600 |
| CD11b | BUV563 | BD Biosciences | M1/70 | 1:4000 | Cat: 741242<br>RRID: AB_2870793 |
| F4/80 | BUV661 | BD Biosciences | T45-2342 | 1:200 | Cat: 750643<br>RRID: AB_2874771 |
| CD64 | BV711 | BioLegend | X54-5/7.1 | 1:100 | Cat: 139311<br>RRID: AB_2563846 |
| MHCII | BUV395 | BD Biosciences | 2G9 | 1:3000 | Cat: 743876<br>RRID: AB_2741827 |
| PDL1 (CD274) | BUV737 | BD Biosciences | M1H5 | 1:100 | Cat: 741877<br>RRID: AB_2871203 |
| PDL2 (CD273) | BV650 | BD Biosciences | T725 | 1:100 | Cat: 740624<br>RRID: AB_2740321 |
| Ly6G | SB550 | BioLegend | 1A8 | 1:1000 | Cat: 127664<br>RRID: AB_2860671 |
| Siglec F | BV480 | BD Biosciences | E50-2440 | 1:400 | Cat: 746668<br>RRID: AB_2743940 |
| Ly6C | PerCP/Cy5.5 | BioLegend | HK1.4 | 1:1000 | Cat: 128012<br>RRID: AB_1659241 |
| CD3 | BV750 | BioLegend | 17A2 | 1:100 | Cat: 100249<br>RRID: AB_2734148 |
| CD98 | BUV615 | BD Biosciences | H202-141 | 1:400 | Cat: 752360<br>RRID: AB_2875877 |

| | | | | | |
|---|---|---|---|---|---|
| CD301 | PE/Dazzle594 | BioLegend | LOM-14 | 1:400 | Cat: 145714<br>RRID: AB_2819896 |
| CD38 | BUV805 | BD Biosciences | ab90 | 1:400 | Cat: 741955<br>RRID: AB_2871263 |

**Table S1:** Antibodies for metabolic and inflammation profiling.
Adapted from (41)
